# Complementary Single-Cell Microflow HILIC and Ion Pair LC-MS Reveal Bystander Metabolic Effects in a Macrophage Model of Tuberculosis

**DOI:** 10.64898/2026.06.22.733771

**Authors:** Abigail Cook, Rahul Deshpande, Abigail E. Ellis, Ryan D. Sheldon, Claire Davison, Jordan Pascoe, Susan Bird, Dany JV Beste, Melanie Bailey

## Abstract

Single-cell metabolomics remains analytically challenging due to the low abundance and chemical diversity of metabolites in individual cells. We have developed complementary microflow HILIC and ion pair LC-MS methods to expand metabolite coverage in single macrophages. Ion pair LC-MS was applied to single cells for the first time, enabling retention of highly polar and ionic metabolites that elute early under conventional reversed-phase conditions. Across *Mycobacterium bovis* BCG infected, uninfected bystander, and control unexposed THP-1 macrophages, both microflow methods detected significantly more features than a previously reported analytical-flow HILIC method. The two microflow methods provided complementary chemical space, together yielding 633 unique named metabolites with MS^2^ spectra. This depth enabled pathway-level interpretation at single-cell resolution, revealing infection-associated changes in purine-, arginine-, glutathione-, and one-carbon folate-associated metabolism. Metabolite-level interrogation indicated shared purine and amino acid changes in both infected and neighbouring macrophages, while revealing a distinct bystander phenotype characterised by elevated glycine and heterogeneous ATP levels. Finally, we demonstrate sequential IP and HILIC analysis of the same single cell, establishing a route toward maximal coverage from individual cells. These results position microflow HILIC and IP LC-MS as powerful, orthogonal strategies for advancing single-cell metabolomics and unveiling heterogeneity within complex biological microenvironments.

**Table of Contents:** Figure made in BioRender.

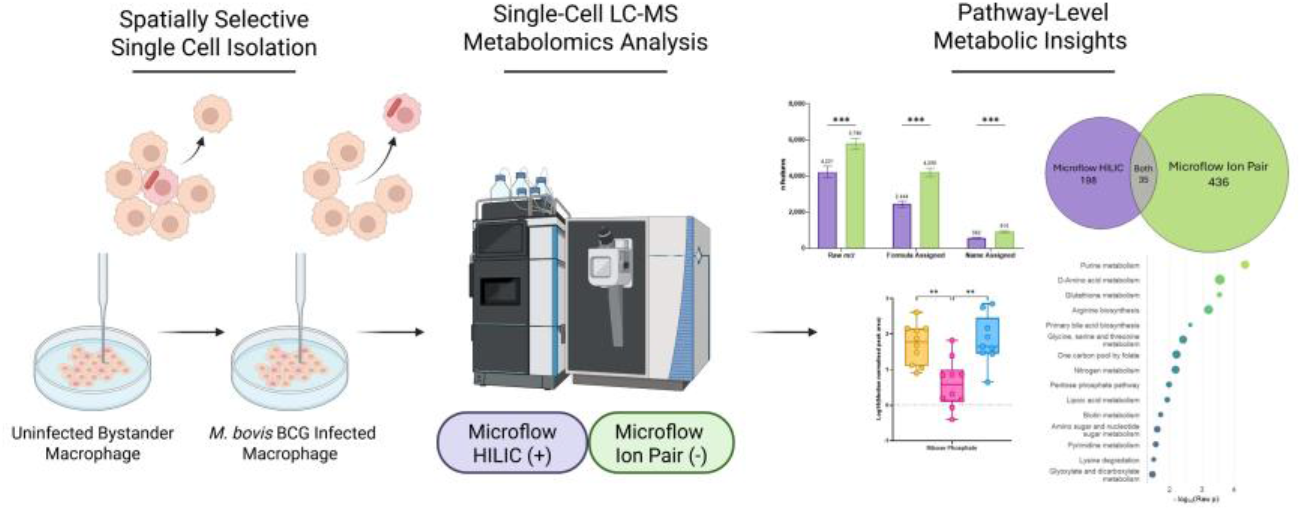

## Introduction

Single-cell metabolomics offers the opportunity to explore cell-to-cell heterogeneity, identify metabolic sub-populations, and to link metabolic phenotypes to infection status, neighbouring cell states or micro-environment.^1,2^ Single-cell metabolomics is a rapidly expanding field, but it faces unique analytical challenges arising from the extreme chemical diversity, dynamic range and ultra-low abundance of metabolites.^3^ These constraints push the limits of current measurement instrumentation and workflows. High-resolution mass spectrometry offers a powerful route to profile low abundance metabolites, since it provides highly sensitive and multiplexed analysis across a broad chemical space. ^4^

Several mass spectrometry-based strategies can perform single-cell metabolomics.^5–8^ Mass spectrometry imaging (MSI) techniques provide rapid, spatially resolved metabolite information at single or even sub-cellular resolution. Techniques such as Secondary Ion Mass Spectrometry (SIMS) and Matrix Assisted Laser Desorption Ionisation-Mass Spectrometry (MALDI-MS) can operate at nanometre and low micrometre pixel sizes respectively, allowing subcellular resolution. However, these approaches cannot analyse cells in their native physiological state, typically requiring chemical fixation or pretreatment with a matrix, or both.^9–11^ Additionally, MSI lacks chromatographic separation prior to ionisation, leaving the measurement susceptible to matrix effects and ion suppression, which limits metabolite coverage. Direct nano-electrospray ionisation (ESI) MS analysis of live single cells offers an alternative, but it suffers from similar challenges to MSI, including the absence of separation and vulnerability to ion suppression.^5,12,13^

Liquid chromatography-mass spectrometry (LC-MS) is widely applied in bulk metabolomics to improve coverage, annotation confidence, and resolve structural isomers in complex biological samples^14^. LC-MS has also been widely adopted in single-cell proteomics, and more recently demonstrated for single-cell lipidomics.^15–21^ However, its development for single-cell metabolomics remains limited.^22–24^ Early LC-MS approaches for profiling metabolites in single cells have shown promise for distinguishing cell populations, but provide limited coverage and limited MS^2^ information.^5,25^ This means accurately annotating analytes, or interpreting biological pathways is difficult. Expanding both coverage and annotation confidence therefore remains a central challenge for single-cell metabolomics.

In this work, we exploit the ability to tune column chemistries and flow rates to separate molecules with a range of different polarities in single cells and thereby enhance coverage of the metabolome. Although reversed-phase (RP) chromatography is ubiquitous in laboratories, the retention and separation of polar and hydrophilic metabolites is extremely difficult without derivatisation. We have previously shown that hydrophilic interaction (HILIC) chromatography can be applied to single cells to detect hydrophilic and some hydrophobic compound classes, including fatty acyls and prenol lipids.^5^ However, microflow LC-MS offers the promise of further gains in sensitivity, because lower flow rates in combination with a smaller probe generate a finer, more efficiently ionised electrospray plume, improving ionisation efficiency, ion sampling, and chromatographic peak shape, ultimately increasing signal-to-noise.^26^ Despite these advantages, microflow HILIC has not previously been reported for single-cell metabolomics.

Ion Pair (IP) chromatography provides an additional, orthogonal strategy for retaining both polar and non-polar analytes on the same column. IP utilises a lipophilic ionic additive to form transient hydrophobic ion pairs, enhancing retention and improving peak shape for analytes that elute early or poorly under standard RP conditions.^27^ IP and HILIC offer complementary selectivity in metabolomics,^28^ yet, to our knowledge, IP-based LC-MS has not previously been applied to single-cell metabolomics.

Here, we develop and integrate low-flow HILIC and IP LC–MS methodologies to increase the number of metabolites from a range of metabolic classes detected from individual cells. Our workflow is compatible with multiple single cell-isolation strategies, including microfluidics and capillary-based sampling. In this study, we use capillary sampling under microscopy, enabling spatially informed selection of cells with defined phenotypes. We demonstrate how this combined methodology provides new insight into bacterial infection at single-cell resolution.

The interaction between host cells and pathogens is central to understanding infectious diseases. Metabolic reprogramming is increasingly recognised as a key regulator of immune responses, particularly within macrophages.^29^ These cells play a key role in fighting infections, such as *Mycobacterium tuberculosis* (Mtb), the causative agent of tuberculosis. In the early stages of infection, only a small fraction of macrophages become infected^30^, resulting in heterogeneous subpopulations containing both infected and uninfected bystander cells.^31^ A fundamental unanswered question is why some macrophages never become infected. We hypothesise that infected macrophages generate a bystander effect that alters the metabolic response of neighbouring uninfected cells, thereby shaping which macrophages ultimately become infected. Single-cell approaches are essential to test this hypothesis, by resolving infected and bystander macrophage states within heterogeneous populations.

We have applied our spatial single-cell analysis workflow to selectively sample infected and uninfected neighbouring cells from a macrophage model of tuberculosis infection using *Mycobacterium bovis* Bacillus Calmette-Guérin (BCG) as a class 2 surrogate for *M. tuberculosis*. We compare the low-flow rate chromatography methods to a previously published analytical flow method,^5^ demonstrating significant improvements in metabolite coverage and annotation confidence in single cells. We further assess the feasibility of pathways enrichment analysis to detect metabolic differences between single *M. bovis* BCG infected THP-1 macrophages and adjacent uninfected bystander macrophages. Finally, we show how the ion pair and HILIC methods can be applied to the same cells, yielding an average of 229 ± 80 metabolites identified by Human Metabolome Database (HMDB) annotation, each supported with MS^2^ spectra, enabling deeper interpretation.

## Experimental

### Chemicals and Reagents

THP-1 and *Mycobacterium bovis* Bacillus Calmette-Guérin (BCG) cells were obtained from ATCC, USA. Roswell Park Memorial Institute (RPMI) 1640 media and fetal bovine serum (FBS) were acquired from Merck, UK and Thermo Fisher Scientific, UK respectively. Phorbol 12-myristate 13-acetate (PMA) was purchased from Sigma-Aldrich, UK. Middlebrook 7H9 medium was acquired from Sigma-Aldrich, UK and supplemented with 5% (v/v) glycerol (Thermo Scientific, UK), 0.2% (v/v) Tween80 (Sigma-Aldrich, UK) and 5% (v/v) Albumin Dextrose Catalase (ADC) (Remel, US). Dulbecco’s Phosphate Buffered Saline (DPBS) with calcium (Ca^2+^) and magnesium (Mg^2+^) was purchased from Gibco, US.

Solvents, including acetonitrile (ACN), water (H_2_O), methanol (MeOH), formic acid (FA) and acetic acid (AA) were Optima LC-MS grade from Fisher Scientific, UK. tributylamine and medronic acid were purchased from Sigma-Aldrich, US and Agilent Technologies, US respectively. Mixed deuterated amino acid standard was purchased from Merck, UK. Credentialed *Escherichia Coli* Cell Extract was purchased from Cambridge Isotope Laboratories, Inc., US.

### THP-1 Macrophage Derivation and Infection

THP-1 culture and differentiation, and BCG growth and infection was as described in Cook *et al*.^5^ Four dishes of PMA induced THP-1 macrophages were infected with BCG at a multiplicity of infection (MOI) of 10:1 and incubated for four hours, before washing and resuspending with 2 mL DPBS with Ca^2+^ and Mg^2+^. For this work, a fluorescent strain of BCG containing an episomal plasmid was used, constitutively expressing monomeric red fluorescent protein mCherry.^32^ This aided microscopic visualisation of the bacteria. Unexposed THP-macrophages were also cultivated.

### Single-Cell Imaging and Sampling

The SS2000 (Yokogawa, Japan) was used to incubate and isolate single THP-1 macrophages. The incubator was set to 37.5 °C with 5% CO_2_ and humidity on. Each macrophage was aspirated into glass capillary tips with an inner diameter of 10 μm using pre-sampling, sampling and post-sampling pressures of 1.80, −17.70 and 0.00 kPa respectively.

Infected cells were identified through detection of mCherry red fluorescence within the macrophage, using excitation and emission wavelengths of 561 nm and 617/73 nm respectively for 500 ms, verified through *z*-stack imaging. Successful aspiration of macrophages into capillaries was confirmed through video footage of the sampling, as well as comparison of *z*-stack imaging before and after aspiration.

For each single cell group (control, infected, bystander), cells were sampled as follows: 40 infected macrophages (infected cells) and 40 uninfected macrophages, (bystander cells) were sampled from the BCG-infected dish; 40 control macrophages were sampled from an unexposed control dish. Immediately after sampling, each tip was placed on dry ice before being stored at −80 °C. DPBS with Ca^2+^ and Mg^2+^ was also sampled from a dish containing no cells and was used as a blank.

### Sample Preparation

The single cells and PBS blanks in capillary tips were thawed before backfilling with 5 μL 1:16.7 mixed deuterated amino acid internal standard in 1:1 H_2_O:ACN + 0.1% FA using GELoader pipette tips. The internal standard was used to confirm that the sample had transferred effectively from the capillary tip to the vial. The contents of the capillary tips were then eluted into QSert autosampler vials (Supelco, UK) using a 10 mL gas syringe (Hamilton, UK) attached to a syringe driver (KD Scientific Incorporated, USA). Finally, 10 μL 1:1 H_2_O:ACN + 0.1% FA was then added to the vial.

Half of the single cell samples and PBS blanks were then frozen at −80 °C and lyophilized (CoolSafe, SCANVAC) for 2 h prior to shipping to Thermo Fisher, San Jose, US where they were reconstituted in 5 μL 10% MeOH in H_2_O. For a sub-set of cells, the reconstitution volume was increased to 10 μL. These cells were then analysed on both low flow rate methods, detailed in **LC-MS Analysis** section.

### LC-MS Analysis

LC-MS analysis was conducted on two different instrumental set ups to assess the coverage of metabolites detected in single macrophages. 20 single cells from each sample group (control, infected, bystander) were analysed using an analytical flow HILIC method at the University of Surrey, UK, described previously^5^. The remaining cells were freeze-dried and shipped to Thermo Fisher, USA, for analysis on microflow IP and HILIC methods.

The microflow IP and HILIC chromatography methods were adapted for single cell analysis at Thermo Fisher USA using a Vanquish Neo Liquid Chromatography System coupled to an Excedion Pro Quadrupole-Orbitrap mass spectrometer. Method adaptations, such as column temperature, gradient, injection volume, reconstitution solvent composition and volume, RF% and voltage. Table 1 compares the key LC-MS parameters for the optimised methods. Further details can be found in Supporting Information Table S1.

**Table 1.** Summary of methods developed to profile metabolites in single cells on both Q Exactive™ Plus and Excedion™ Pro.

|  | Analytical Flow HILIC | Microflow HILIC | Microflow IP |
| --- | --- | --- | --- |
| Chromatography | Hydrophilic | Hydrophilic | Ion Pair |
| Mobile Phases | A: H<sub>2</sub>O + 0.1% FA<br>B: ACN + 0.1% FA | A: H<sub>2</sub>O + 0.1% FA<br>B: ACN + 0.1% FA | A: H<sub>2</sub>O + 3% MeOH + 15 mM AA + 10 mM TBA + 1 μM MA<br>B: MeOH + 15 mM AA + 10 mM TBA + 1 μM MA |
| Flow rate (μL/min) | 400 | 5 | 1 |
| Gradient Time (min) | 15 | 26 | 20 |
| Injection volume (μL) | 15 | 5 | 5 |
| Polarity | Positive (+) | Positive (+) | Negative (-) |
| MS Order | Full MS | MS<sup>2</sup> DDA | MS<sup>2</sup> DDA |
| Resolution at 200 *m/z* (MS<sup>1</sup>/ MS<sup>2</sup>) | 140,000 | 120,000/15,000 | 120,000/15,000 |

The ion pair method, adapted from House *et al*.^33^ and developed by the Van Andel Institute Mass Spectrometry Core (RRID:SCR_024903), utilised an EasySpray PepMap Neo C18 phase column (75 μm x 150 mm) (Thermo Fisher Scientific, USA) heated to 60 °C. Mobile phases A and B were H_2_O + 3% Methanol (MeOH) + 15 mM acetic acid (AA) + 10 mM tributylamine (TBA) + 1 µM medronic acid (MA) and MeOH + 15 mM AA + 10 mM TBA + 1 µM MA, respectively. IP gradient is described in Supplementary Information Table S2.

The HILIC method reported in Cook *et al*.^5^ was adapted for use on the Vanquish Neo. Method adaptations included flow rate, gradient and column temperature. An amide phase column (3 x 150 mm) (Waters, UK) was heated to 30 °C. Mobile phase A was H_2_O + 0.1% FA and B was ACN + 0.1% FA. Microflow HILIC gradient is described in Supplementary Information Table S3.

### Data Analysis

All mass spectrometry datasets were processed using CompoundDiscoverer 3.4 (Thermo Scientific, USA). The Human Metabolome Database (HMDB 5.0, https://hmdb.ca/)^34^ was used as a reference library. RStudio 2026.01.0+392 (R 4.3, Posit, USA, http://www.posit.co/), Prism 10 (GraphPad Software Incorporated, USA) and MetaboAnalyst 6.0 (https://www.metaboanalyst.ca/home.xhtml)^35^ were used for data handling, statistical testing, and figure generation. Metabolite classes were determined by using ClassyFire Batch by Fiehn Lab (https://cfb.fiehnlab.ucdavis.edu/#!#%2F), based on the application developed by the Wishart Research Group.^36^ Additionally, RStudio, TraceFinder 5.2 (ThermoScientific, USA) and MetaboAnalyst 6.0 were employed for pathway enrichment analysis, utilizing the KEGG metabolic pathways database (Nov 2025), and ensuring >10% pathway coverage. Images were processed using FIJI (ImageJ 1.54r, https://imagej.net/ij/).^37^

Data analysed within MetaboAnalyst was log10 transformed and median normalised for univariate analysis, multivariate analysis data was also autoscaled, after features with >80% missing values were removed and remaining missing values were imputed with 1/5 of the minimum positive value per feature.

All single cell data was blank corrected to capillary sampled PBS with Ca^2+^ and Mg^2+^. For the cells that were split between microflow IP and HILIC methods, PBS blanks were also split and analysed accordingly.

## Results and Discussion

Isolation of single cells was completed using SS2000 capillary sampling, giving us the ability to identify infected cells microscopically and selectively aspirate cells in their native state. Sampling of an infected (Cell A) and uninfected bystander (Cell B) cell into glass capillary tips (black dot) is illustrated in Figure 1. The bystander cell group consisted of uninfected cells from macrophage cultures dosed with mCherry BCG, confirmed by the absence of red-fluorescent bacteria.

**Figure 1.**
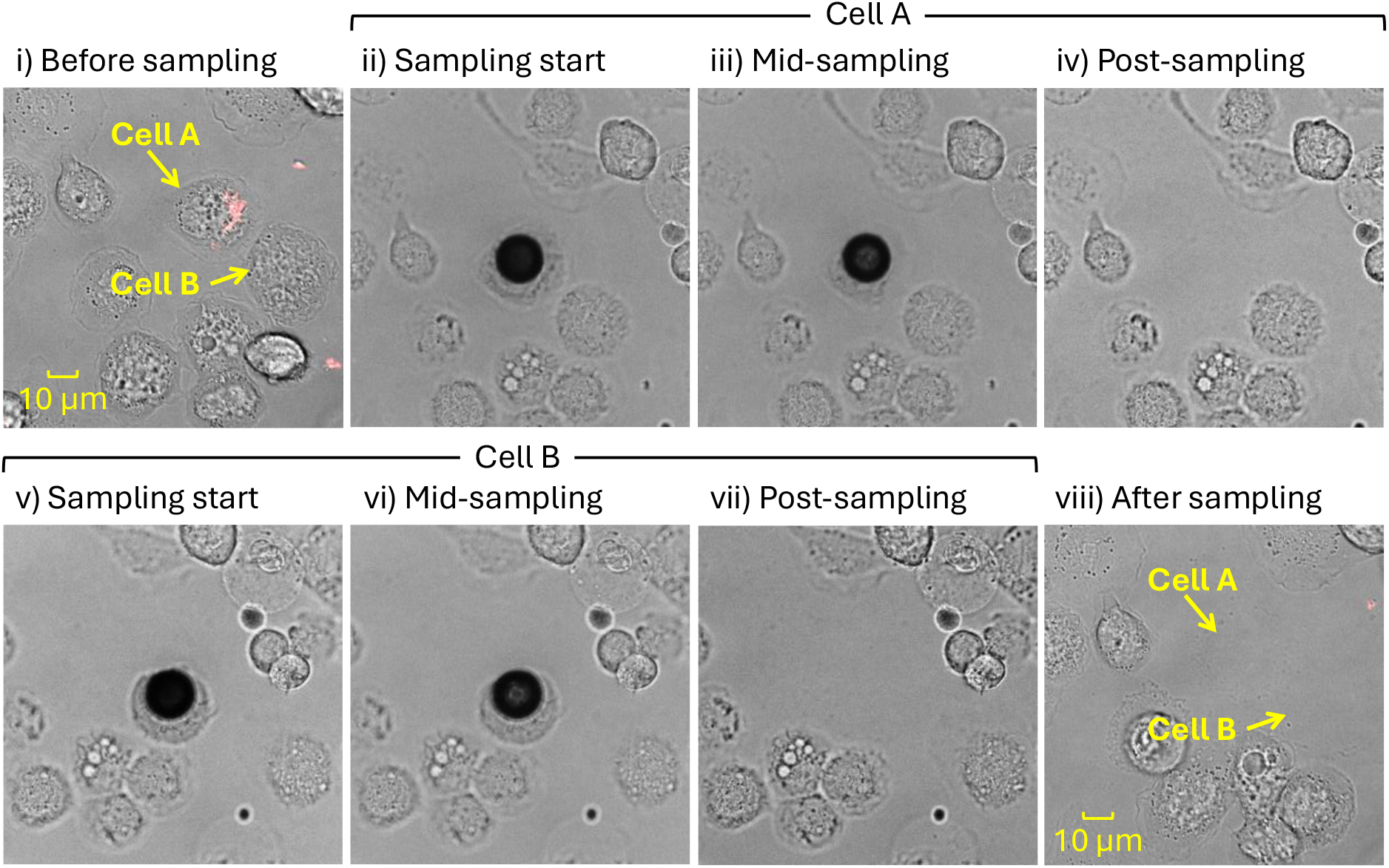
Sampling of infected single cell (A) containing red fluorescent mCherry BCG and uninfected bystander single cell (B). i) Image take prior to sampling cells A and B. ii-iv) sampling of infected cell A before, during and after sampling where the black circle is the glass capillary the cell is sampled into. v-vii) sampling progress of uninfected bystander Cell B. viii) Image taken after sampling cells A and B. 40x brightfield and fluorescent (Ex 561 nm, Em 617/73 nm for 500 ms) images have been stacked in i) and viii) with 10 μm scale bar.

Freeze drying single cells opens the opportunity to ship cells between laboratories, which can be helpful if the cell picking expertise and equipment is not co-located with high end mass spectrometry. Although freeze drying has been explored for single cell lipid analysis, its impact on hydrophilic metabolites in single cells deposited in LC-MS vials has not, to our knowledge, been assessed.^20^ We therefore explored whether freeze drying alters the metabolite profiles of single cells. Across single macrophages, we observed no significant difference in the average number of features detected with/without freeze drying (SI Figure S1). While the number of formula-level annotations differed (p = 0.045), there was no significance in the number of name-annotated features, and most annotations overlapped between conditions (SI Figure S2). Median peak areas were also unchanged (SI Figure S3). These results demonstrate that freeze-dying does not measurably affect single macrophage metabolite profiles, supporting its use for this interlaboratory study.

The microflow ion-pair (IP) method was optimised to detect metabolites at the single-cell level. Supplementary Table S4 lists analytes monitored during method optimisation, covering a range of metabolite classes, such as amino acids, nucleosides, coenzymes, sugars and other hydrophilic species. Initial optimisation focused on improving sensitivity by varying the voltage and RF lens percentage, as shown in Supplementary Figures S4 and S5, respectively. Increasing the magnitude of negative electrospray voltage generally increased in median-normalised peak intensity, whereas the effect of RF lens percentage was more metabolite dependent. An RF lens value of 75% was therefore selected as a suitable compromise across the metabolite panel.

Sample solvent composition is known to influence chromatographic performance, including retention, peak shape and signal intensity, critical for sensitive and robust LC-MS.^38,39^ Supplementary Figure S6 shows extracted ion chromatograms demonstrating the effect of increasing methanol (MeOH) content of the sample solvent from 5-20% in 5% increments and 20-40% in 10% increments. Though some analytes, such as glutathione and coenzyme A, showed optimal intensity at 15% MeOH, the peak shape deteriorated for other analytes, particularly those that elute early or late in the gradient. Therefore, 5% MeOH was chosen to maintain acceptable chromatographic performance across the analyte panel.

Once both microflow (IP and HILIC) methods were optimised, we analysed all single macrophages sampled by the SS2000 and compared the coverage of metabolites to the analytical flow HILIC method. Figure 2A compares the number of features assigned by *m/z* value, formula and compound name detected by the different methods, showing the microflow methods detect significantly more features. Additionally, more features were detected using IP chromatography than by either HILIC method. This reflects the ability of ion-pairing to extend reversed-phase retention to highly polar and ionic metabolites that would otherwise elute early, whereas HILIC selectively retains hydrophilic species. Interestingly, from the full scan data, there was no significant difference in the number of name-annotated features detected between the two HILIC methods. When assessing the classes of these named features, as shown in Figure 2B, there is similarity between the outputs of both HILIC methods contain similar proportions of compounds in each of the top 15 classes.

**Figure 2.**
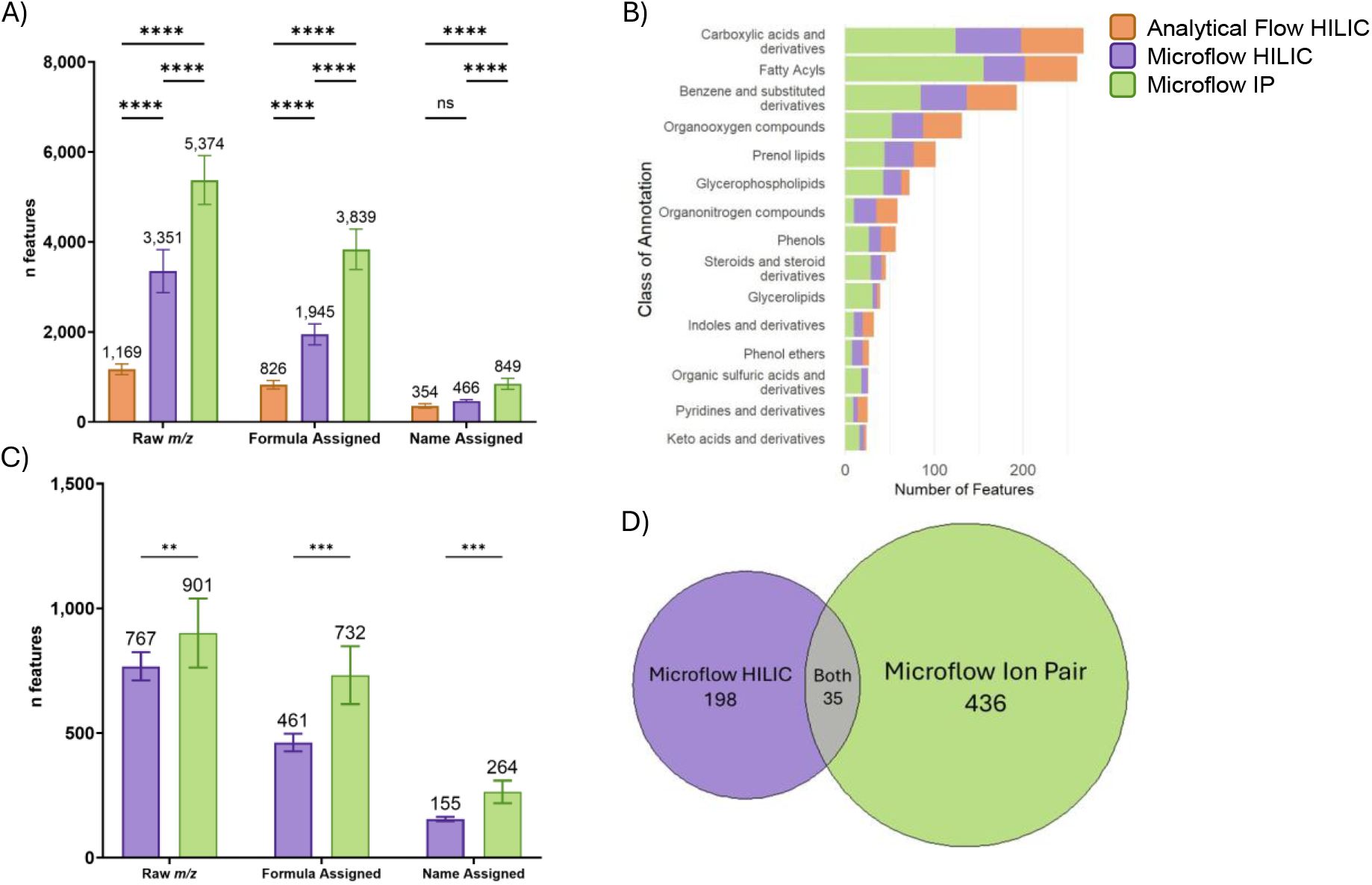
Comparison of features detected in single cells analysed by analytical flow HILIC (orange, n = 60), microflow HILIC Excedion Pro (purple, n = 16) and microflow IP Excedion Pro (green, n = 29) methods. A) Average features (±SD) detected within single cells by each analysis method. Significance was calculated by a 2-way ANOVA with Holm-Šídák correction for multiple comparisons. B) Distribution of name annotated features in the top 15 classes per method. C) Average features with MS^2^ spectra (±SD) detected within single cells analysed by both microflow methods, excluding the analytical flow method which was run in full scan mode only, due to the slow duty cycle of the mass spectrometer. Mann-Witney U tests with Holm-Šídák correction for multiple comparisons were performed to determine significance. D) Venn diagram displaying the overlap of all unique named annotations with MS^2^ spectra for both microflow methods.

Figure 2C shows an additional advantage of the microflow rate methods in collecting MS^2^ spectra. The IP method detected more (Mann-Witney U test) MS^2^ features per cell than the microflow HILIC method. The analytical flow method was run in full scan mode only due to the slow duty cycle of the Q Exactive, so was not compared. Figure 2D shows the total number of named metabolite identifications with MS^2^ spectra and the subsequent limited overlap between the IP and HILIC microflow methods, which were run in negative and positive ionisation mode respectively. Together, these methods cover an unprecedented array of polar and non-polar metabolite classes in single cells.

To explore differences in metabolic features between control, uninfected and bystander cells, multivariate analysis was performed for all *m/z* features, as presented in Figure 3. Both the HILIC and IP method showed clear differences in metabolite intensity between control and infected cells, as evidenced by the PLS-DA (Figure 3A and D) and the heatmaps (Figure 3B,C and E,F). Leave one out cross validation (LOOCV) analysis for both microflow IP and HILIC PLS-DA plots can be found in SI Figure S7 and S8. Interestingly, in both methods, the heatmaps in Figure 3B,C and E,F show that the bystander cells had greater similarity to the infected cells than the unexposed controls. There are also two bands of features that differentiate the bystanders from the infected cells, information that cannot be obtained through bulk analysis.

**Figure 3.**
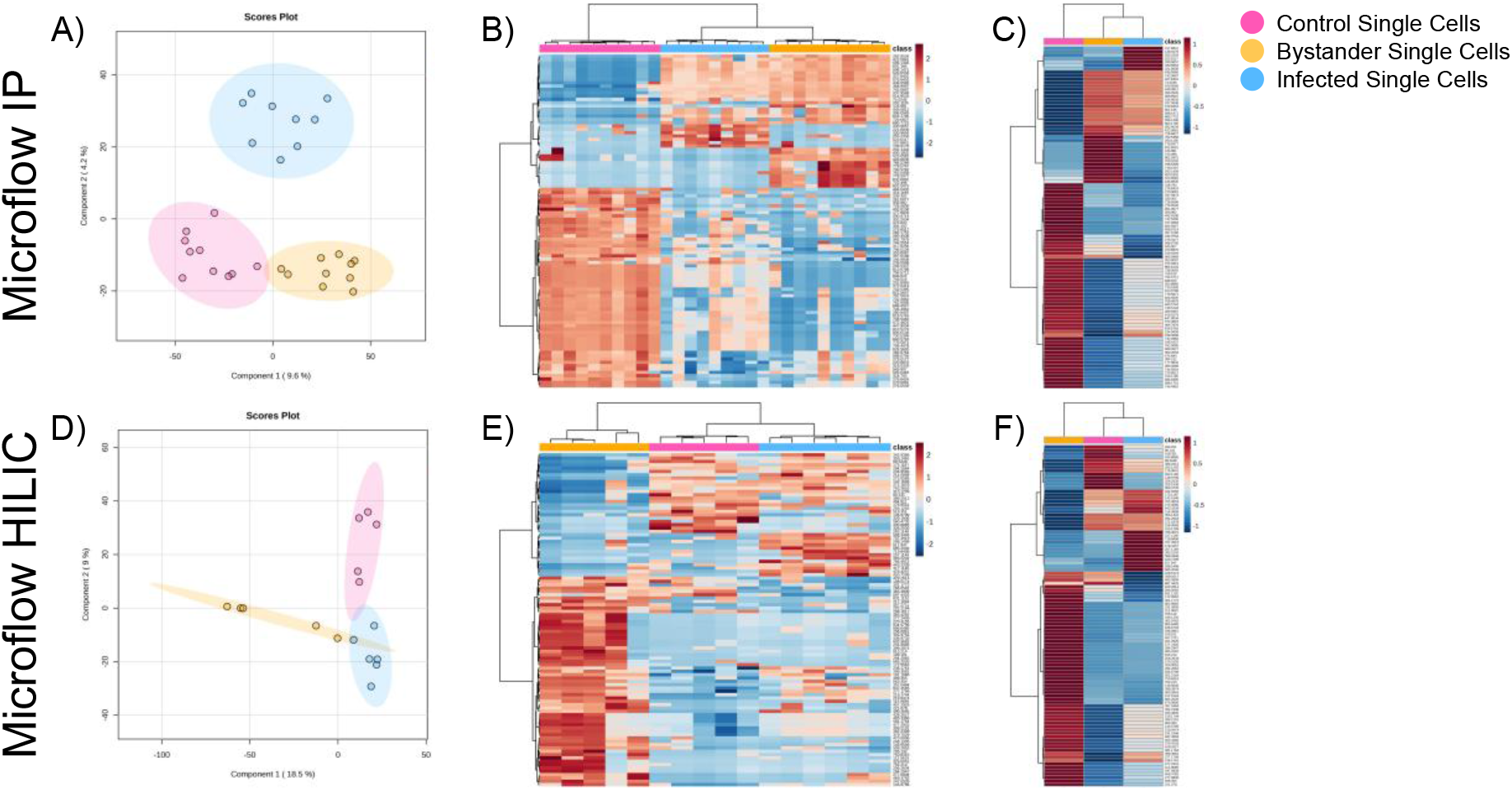
Multivariate analysis of *m/z* features within unexposed control (pink), uninfected bystander (yellow) and BCG infected (blue) single macrophages using microflow IP (n = 29) and HILIC (n = 16) methods. A) and D) are PLS-DA of IP and HILIC Excedion methods respectively. Heatmaps displayed in B) and E) display the top 100 significant *m/z* features by ANOVA comparing the three biological groups for IP and HILIC Excedion methods respectively, with figures C) and F) summarising the average values.

Pathway-level interpretation has never been performed at a single cell level due to limited metabolite coverage and identification confidence. The detection of a total of 633 unique metabolites with MS^2^ spectra in the dataset now opens the possibility to perform a more in-depth analysis of the data, which is explored here. We first applied the human metabolite database (HMDB) to ensure a fully human database for feature annotation which were then translated to KEGG IDs for pairwise pathway enrichment analysis for each sample group within each method. As in bulk analysis, a given *m/z* feature may correspond to multiple isomeric metabolites, such as *D/L* or *E/Z* stereoisomers, which cannot be confidently distinguished by accurate mass, even if MS^2^ spectra are recorded. This results in restricting each feature to a single KEGG ID annotation which corresponds to only one of these isomers, excluding biologically plausible pathway members and reducing enrichment sensitivity. To address this limitation, we selected the KEGG IDs relevant to the top 20 KEGG pathways in our dataset and applied targeted analysis using TraceFinder software. This supervised analysis resulted in wider pathway coverage that avoids bias toward a pre-defined metabolite list, improving the confidence in the pathway enrichment analysis. The pairwise results of the enrichment analysis using MetaboAnalyst are in SI Tables S5-10 and the top 15 pathways are summarised in Figure 4.

**Figure 4.**
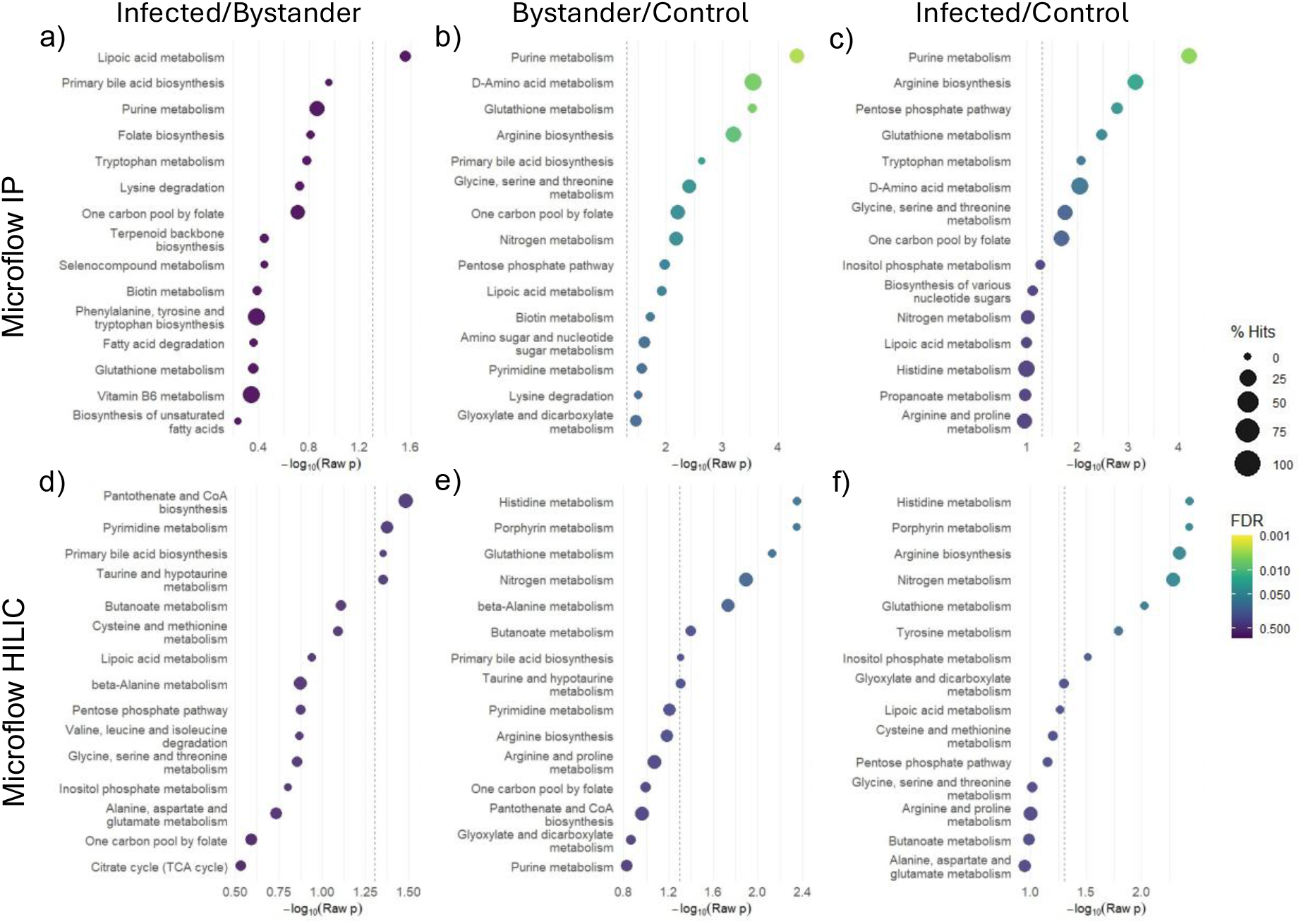
Targeted pairwise quantitative enrichment analysis of BCG infected, uninfected bystander and control unexposed single cells analysed by IP Excedion (A, B, C) or HILIC Excedion (D, E, F). Top 15 pathways (raw *p*-value) for Infected/Bystander, Bystander/Control and Infected/Control pairwise enrichment. Dotted line represents raw *p*-value significance threshold at −log10(0.05). Size of each dot correlates to the percentage of features detected within each pathway and the colour is scaled to FDR significance.

Plotting both raw *p*-value and false discovery rate (FDR) to preserve sensitivity to modest enrichment trends whilst indicating statistical robustness showed that both methods detected metabolite enrichment in several metabolic pathways (Figure 4). In many cases, 25-50% coverage of the pathways was observed, comparable to many bulk analyses, with 77% of the purine metabolism pathway being detected (SI Figure S9). Whilst there was some overlap in enriched pathways between the IP and HILIC methods (e.g. arginine biosynthesis, nitrogen metabolism), pathway coverage varied and some pathways were only identified by a single method. This demonstrates their complementarity and highlights the value of combining both approaches.

Pathway analysis revealed that in both infected and bystander macrophages there were changes in metabolites derived from purine metabolism, arginine biosynthesis, glutathione metabolism, one carbon/folate pathways, relative to unexposed control macrophages (Figure 4B, C, E and F). These pathways are hallmarks of cytokine-driven macrophage activation, observed during Mtb infection.^40–46^ As seen in Figure 4A and D, there are only minor differences in enriched pathways between infected cells and their neighbouring uninfected cells, regardless of analysis method. This indicates that, despite being uninfected, the cells are being influenced by its neighbouring infected cell, suggesting a bystander effect. In this study, we did not track the infection status of the sampled cells over time, so we cannot determine whether these bystander cells were previously infected or are truly never-infected cells. This could be addressed in future work by performing time course imaging in the instrument prior to cell sampling. We hypothesise that this metabolic convergence is driven by signalling factors released by infected cells (e.g. cytokines, chemokines, or microbial products), that protect against infection.

To support the pathway-level analysis (Figure 4), we selected five metabolites detected by the microflow IP method that link the most significantly enriched pathways. Box plots of each feature are shown in Figure 5, with each overlaid dot representing the measurement in a single cell. A ribose phosphate feature consistent with ribose 5-phosphate (Figure 5A), hypoxanthine (Figure 5B) and ornithine (Figure 5C) were significantly enriched in both infected and bystander macrophages relative to controls. These metabolites link some of the enriched pathways, with ribose phosphate connecting the pentose phosphate pathway and purine metabolism, hypoxanthine reflecting purine salvage, and ornithine indicating altered arginine biosynthesis and D-amino acid metabolism. This data further supports our pathway analysis and indicates that BCG induces a bystander effect.

**Figure 5.**
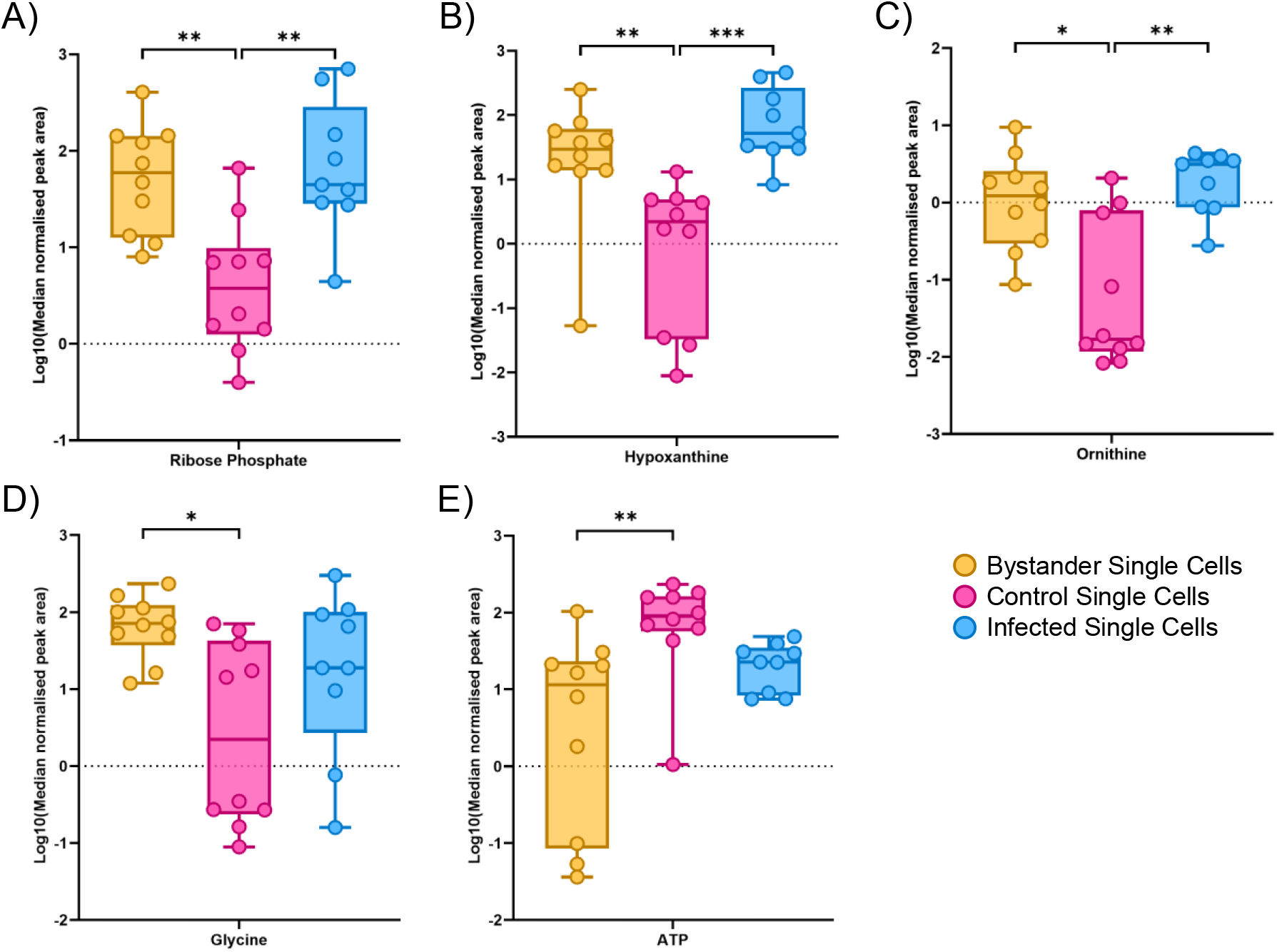
Box and whisker plots displaying the distribution of log10 transformed and median normalised peak areas of BCG infected (blue, n = 9), uninfected bystander (yellow, n = 10) and unexposed control (pink, n = 10) single cells detected using microflow IP method. Box extends to the 25th and 75th percentiles, line in the middle of the box is the median and the whiskers display the minimum and maximum values. Overlaid dots represent the measurement for each single cell. Metabolites displayed are A) Ribose Phosphate, B) Hypoxanthine, C) Ornithine, D) Glycine, and E) Adenosine Triphosphate (ATP). Significance was determined by a Kruskal-Wallis test with Dunn’s correction for multiple comparisons, not significant values are not displayed.

Although the infected and bystander population averages appeared broadly similar, bystander single cells displayed additional features that distinguished them from both infected and unexposed control macrophages. Glycine (Figure 5D) was significantly enriched in the bystander cells when compared to the unexposed control macrophages while adenosine triphosphate (ATP) (Figure 5E) was significantly depleted. A noticeable difference in the distribution of the abundance of glycine and ATP was detected, (see SI Table S11 andS12 for statistical tests). Glycine was found to be less variable in bystander macrophages than in infected or control cells. This suggests bystander macrophages may shift their levels of glycine distinctly in response to exposure to BCG infection.

Conversely, ATP (Figure 5E) was highly variable between bystander macrophages, specifically compared to the infected macrophages, which showed tighter dispersion of ATP. Additionally, the ATP abundance was significantly depleted in bystander macrophages relative to the controls (Kruskal-Wallis test). Mtb infecton is known to reduce ATP-levels.^47^ Here we extend this observation to show this also occurs in uninfected bystander macrophages.

Having demonstrated the utility of applying HILIC and IP separately to different groups of single cells, we next tested the feasibility of performing both methods in sequence on the same cell, with the aim of maximising analyte coverage from a single cell. The methods were unchanged, except that the cells were reconstituted in double volume of injection solvent (10 μL 10% MeOH in H_2_O). The cells were analysed on IP method first followed by the HILIC method the following day.

Figure 6A displays the average features detected by applying the methods sequentially to the same single cell. SI Figure S10 compares the average features in whole cells and half cells, in which there is no significant difference in the number of named features, implying no loss in sensitivity, despite the higher dilution. This is reflected in the number of features with MS^2^ spectra (Figure 6B). Figure 6C shows a PLS-DA of infected, bystander and unexposed control single cells analysed using this sequential method. The PLS-DA shows clear separation between groups, although this may be overfitting due to the lower sample size, reflected in the negative Q2 scores (SI Figure S11). Interestingly, in spite of the smaller sample size (n = 4 cells per group) the heatmaps (Figure 6D and 6E) recapitulate the relationships observed in Figure 3B, C, E and F – specifically, greater similarity between the bystander and infected cells than the controls, and a panel of features that differentiate bystander and infected cells.

**Figure 6.**
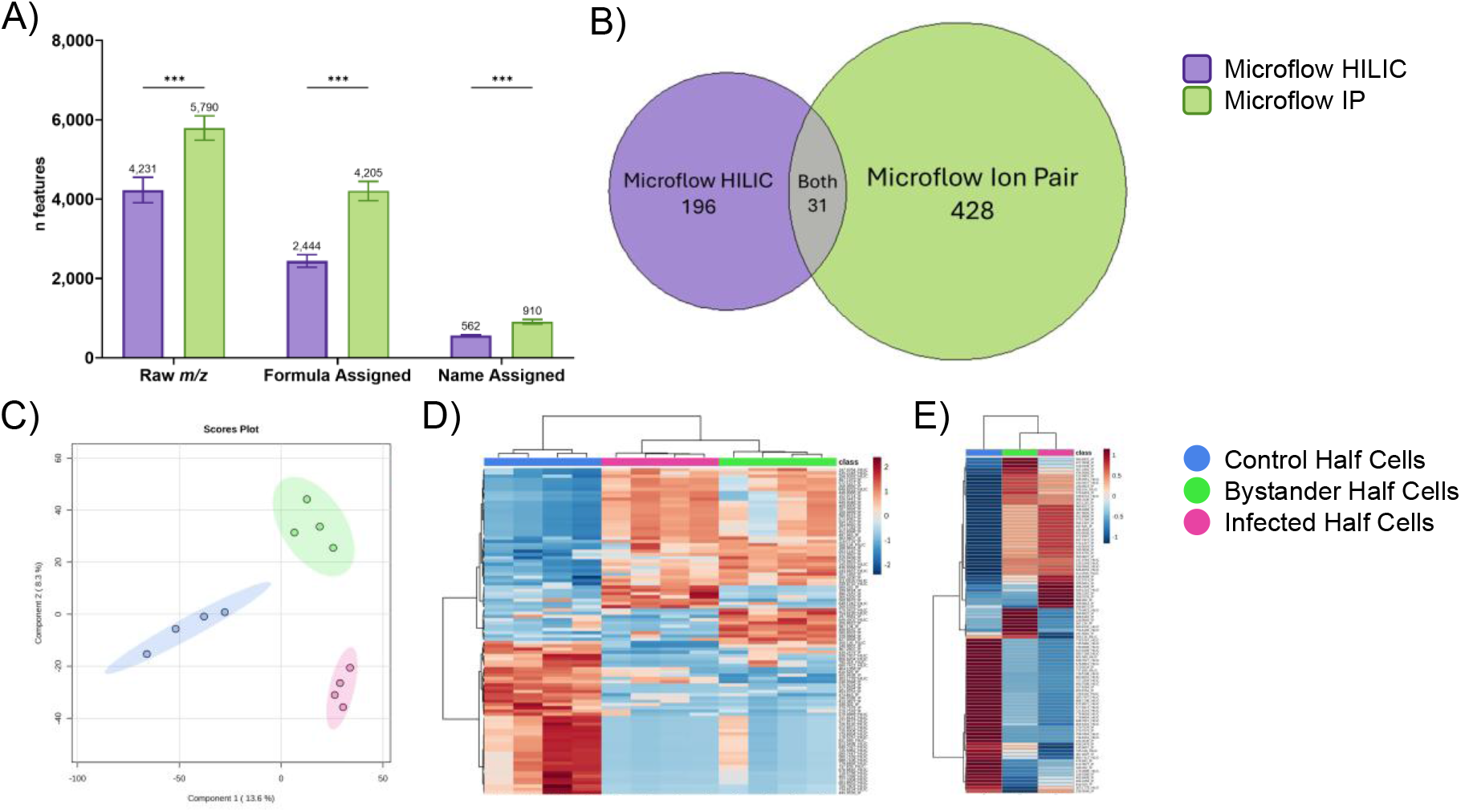
Single cells (n = 12) analysed sequentially by IP (green) and HILIC (purple). A) displays average MS^1^ and MS^2^ features (±SD) detected from single cells split between the two analysis methods. A Mann-Witney t-test with Holm-Šídák correction for multiple comparisons was performed to determine significance. B) Venn diagram displaying the overlap of all unique named annotations with MS^2^ spectra. Figures C, D and E describe features within unexposed control (blue), uninfected bystander (green) and BCG infected (pink) single macrophages, C) PLS-DA of all features detected from both methods. D) Heatmap of top 100 significant (ANOVA) features detected in each single cell. E) Heatmap averaged for each cell group.

By performing sequential microflow IP and HILIC analyses on “half a cell” at a time without loss of sensitivity, we have demonstrated the sensitivity and flexibility of low-flow LC–MS for single-cell metabolomics. Importantly, this opens the possibility of extending LC–MS to subcellular structures. In principle, nuclei, cytoplasmic fractions, or spatially defined subregions aspirated by capillary sampling could be profiled using the same low-flow LC–MS methods, an avenue we plan to explore in future work.

Though the ion pair method provided the most comprehensive metabolite coverage in this study, it has several practical requirements which may limit its accessibility to many laboratories. The use of TBA as an ion pair reagent can cause persistent LC contamination and carryover, often requiring a dedicated instrument to avoid interference with other workflows. Additionally, ion pair reagents supress ionisation of positively charged analytes, reducing the coverage of the metabolites which preferentially ionise in positive mode.^48^ Furthermore, further purification of TBA is required to reduce background interference and achieve the sensitivity required for microflow single-cell analysis. Therefore, while highly effective for polar and anionic metabolites, microflow IP LC-MS remains technically demanding and benefits from complementary LC-MS approaches, such as HILIC-LC-MS.

A further consideration is that the analytical flow and microflow datasets were acquired on different mass spectrometers. As such, the comparison between analytical-flow HILIC and the microflow IP/HILIC methods is not a strict instrument-controlled comparison. Differences in ion optics, duty cycle, and MS^2^ acquisition strategies inevitably contribute to the higher feature counts observed in the microflow datasets. Nevertheless, the chromatographic trends remain informative: both microflow methods consistently increased the number of detected features, improved MS^2^ coverage, and expanded the chemical space accessible from single cells. These gains reflect the combined effects of lower flow rates, improved ionisation efficiency, and orthogonal chromatographic selectivity, rather than instrument performance alone. Thus, while absolute feature numbers cannot be directly attributed to chromatography in isolation, the overall findings robustly demonstrate the value of microflow HILIC and ion-pair LC-MS for enhancing single-cell metabolomics.

A remaining limitation of the current workflow is its relatively low analytical throughput. The combination of single-cell sampling, low-flow chromatography, and MS^2^ acquisition provides exceptional depth, but results in long analysis times per-cell (20-26 min per cell). Increasing throughput will be essential for scaling single-cell metabolomics to larger populations or time-course studies. Future work could explore strategies such as faster microflow gradients, parallelised LC channels, higher-speed MS^2^ acquisition to increase the number of cells analysed per unit time. Furthermore, a greater number and quality of MS^2^ spectra and analysis of metabolite standards would further improve the confidence in metabolite annotation and identification. Importantly, any gains in throughput must preserve the sensitivity and MS^2^ coverage that underpin the pathway-level insights demonstrated here. Developing higher-throughput versions of these methods would enable more comprehensive mapping of infection-associated metabolic states across larger and more diverse single-cell populations.

## Conclusions

In this work, we have established microflow ion pair and hydrophilic LC-MS as complementary strategies for spatial single-cell metabolomics. IP chromatography is applied to single cells for the first time, extending reversed-phase retention to highly polar and ionic metabolites and substantially increasing feature detection and MS^2^ coverage. Together, the two microflow methods provide broad and orthogonal chemical space, enabling the identification of 633 unique metabolites with MS^2^ spectra and supporting pathway-level interpretation at single-cell resolution. Applying these methods to BCG-infected macrophages revealed similarities and differences between infected and bystander cells, highlighting infection-associated rewiring of purine, arginine, glutathione, and one-carbon by folate pathways. Further probing into metabolic features driving the pathway enrichments observed revealed a shared metabolic response in infected and bystander macrophages, while also uncovering distinct bystander-associated phenotype characterised by the coordinated glycine enrichment and heterogeneous energetic behaviour. Future single-cell metabolomic studies investigating bystander macrophage heterogeneity over time will reveal cell-to-cell signalling mechanisms that prime uninfected cells and shape their subsequent responses to infection. We have also demonstrated that sequential IP and HILIC analysis can be performed on the same single cell without significant loss of sensitivity, offering a route to maximise metabolite coverage from limited material. These advances position microflow LC-MS as a powerful platform for spatial single cell metabolomics and open new opportunities for mechanistic studies of host-pathogen interactions.

## Author Contributions

M.B., D.J.V.B., S.B., R.D. and A.C. conceived and designed the study. A.C. carried out all cell culture, with assistance from J.P., sampled all single cells and prepared all samples. A.E.E. and R.D.S. developed the microflow IP LC-MS method. A.C. and R.D. carried out microflow LC-MS optimisation and analysis. A.C. and C.D. conducted analytical flow LC-MS. A.C. analysed all data and wrote the article with M.B. and D.J.V.B. All authors contributed to the article and approved the final version submitted.

## Supporting Information

Additional tables and figures including details of all LC-MS methods, effect of freeze drying on metabolic features in single cells, method optimisation data, chromatograms, lists of pathways enriched, KEGG purine pathway map, and supporting statistics.

## Acknowledgements

The authors of this publication would like to thank Thermo Fisher Scientific for providing facilities and consumables to conduct the microflow analyses. An acknowledgement also goes to Yokogawa Corporation for their technical support with the SS2000 Single Cellome. Finally, thanks go to Harpreet Atwal and the FHMS technical staff at the University of Surrey for their technical support.

