## Supplementary Information - Figures and Tables for "Complementary Single-Cell Microflow HILIC and Ion Pair LC-MS Reveal Bystander Metabolic Effects in a Macrophage Model of Tuberculosis"

Table of Figures

Table of Tables

Table S1. Summary of all LC-MS parameters for analytical flow HILIC, microflow HILIC and microflow IP methods.

|  | Analytical Flow HILIC | Microflow HILIC | Microflow IP |
| --- | --- | --- | --- |
| Chromatography | HILIC | HILIC | Ion Pair |
| Column | BEH amide, 2.1 x 150 mm, 1.7 μm | BEH amide, 3 x 150 mm | PepMap Neo C18, 75 μm x 150 mm, 2 μm |
| Flow rate (μL/min) | 400 | 5 | 1 |
| Gradient Time (min) | 15 | 26 | 20 |
| Injection volume (μL) | 15 | 5 | 5 |
| Source | HESI | EasySpray + 15 um bullet-type emitter | EasySpray + Integrated column and emitter |
| Polarity | Positive (+) | Positive (+) | Negative (-) |
| MS Order | Full MS | MS^2^ DDA | MS^2^ DDA |
| Resolution at 200 *m/z* (MS^1^/ MS^2^) | 140,000 | 120,000/15,000 | 120,000/15,000 |
| Mobile Phases | A: H2O + 0.1% FA  B: ACN + 0.1% FA | A: H2O + 0.1% FA  B: ACN + 0.1% FA | A: H2O + 3% MeOH + 15 mM AA + 10 mM TBA + 1 µM MA  B: MeOH + 15 mM AA + 10 mM TBA +1 µM MA |
| Mobile Phases | A: Water + 0.1% Formic Acid  B: Acetonitrile + 0.1% Formic Acid | A: Water + 0.1% Formic Acid  B: Acetonitrile + 0.1% Formic Acid | A: Water + 10 mM TBA + 0.01% Acetic Acid  B: Methanol + 10 mM TBA + 0.01% Acetic Acid |
| AGC Target  (MS^1^/ MS^2^) | 1e6 | 1e6/1e5 | 1e6/1e5 |
| Max IT | 256 ms | Auto | Auto |
| Scan range (*m/z*) | 50-750 | 67-850 | 67-850 |
| Voltage (V) | 4000 | 2500 | 2000 |
| Ion Transfer Tube Temperature (°C) | 275 | 350 | 350 |
| RF Lens | 50 a.u | 75% | 75% |
| Collision Energy (%) | N/A | 35 | 35 |
| Number of Dependent Scans | N/A | 5 | 5 |
| Dynamic Exclusion Filter | N/A | Exclude after n = 1 time for 3 s | Exclude after n = 1 time for 3 s |

Table S2. Microflow IP gradient.

| Time (min) | Flow rate (μL/min) | Mobile Phase B (%) |
| --- | --- | --- |
| 0.0 | 5 | 5 |
| 5.0 | 5 | 5 |
| 5.5 | 1 | 5 |
| 6.0 | 1 | 5 |
| 8.0 | 1 | 20 |
| 10.0 | 1 | 46 |
| 13.0 | 1 | 99 |
| 16.0 | 1 | 99 |
| 17.0 | 1 | 5 |
| 20.0 | 5 | 5 |

Table S3. Microflow HILIC gradient.

| Time (min) | Flow rate (μL/min) | Mobile Phase B (%) |
| --- | --- | --- |
| 0.0 | 5 | 100 |
| 5.0 | 5 | 100 |
| 13.0 | 5 | 0 |
| 18.0 | 5 | 0 |
| 18.1 | 5 | 100 |
| 26.0 | 5 | 100 |

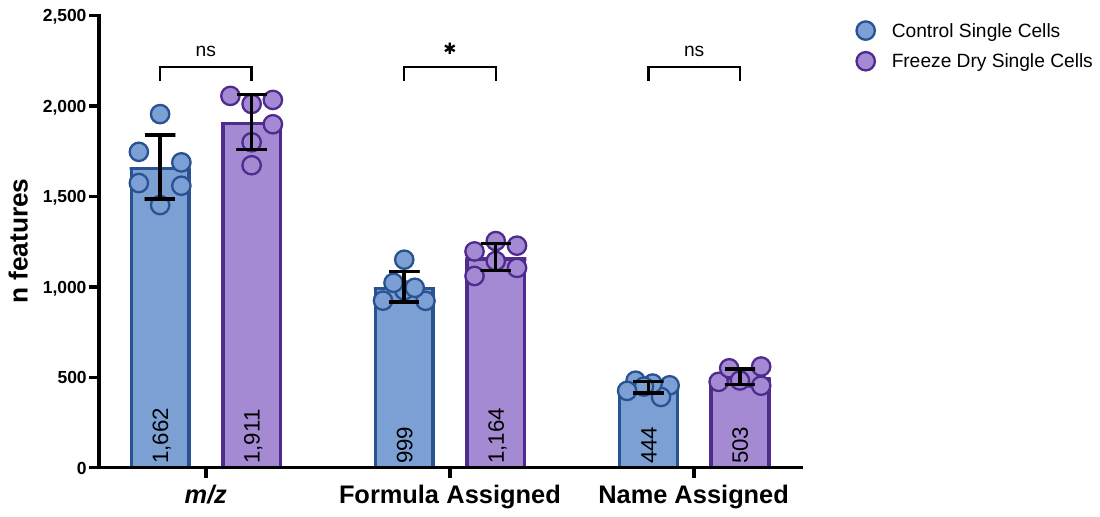

Figure S1. Average features (±SD) detected within freeze dried (blue) and control (purple) single cells by HILIC Q Exactive Plus method. Significance was calculated by a Mann-Witney U t-tests with Holm-Šídák correction for multiple comparisons were performed to determine significance.

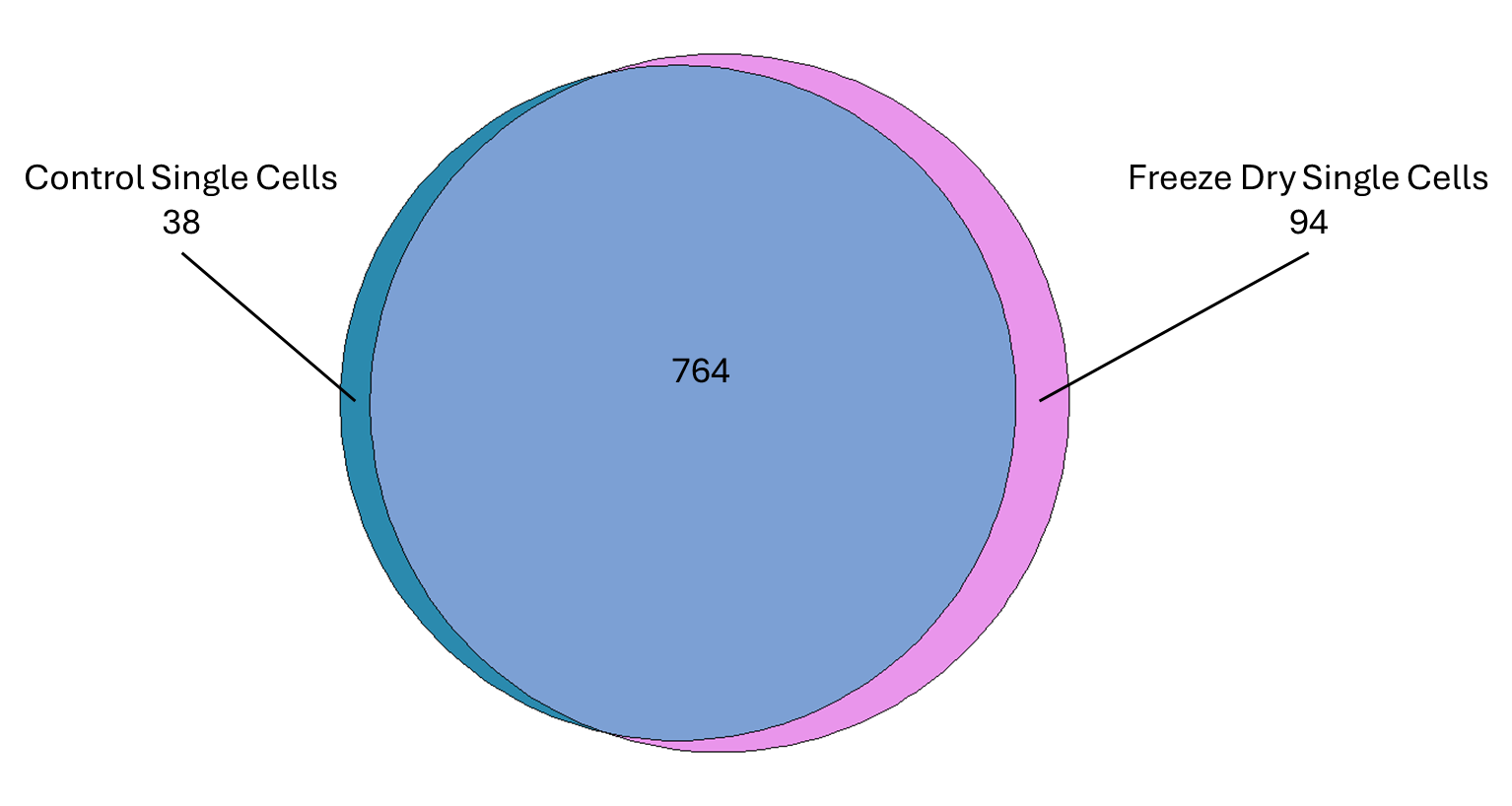

Figure S2. Overlap of all named annotations between freeze dried and control single cells analysed using HILIC Q Exactive Plus method.

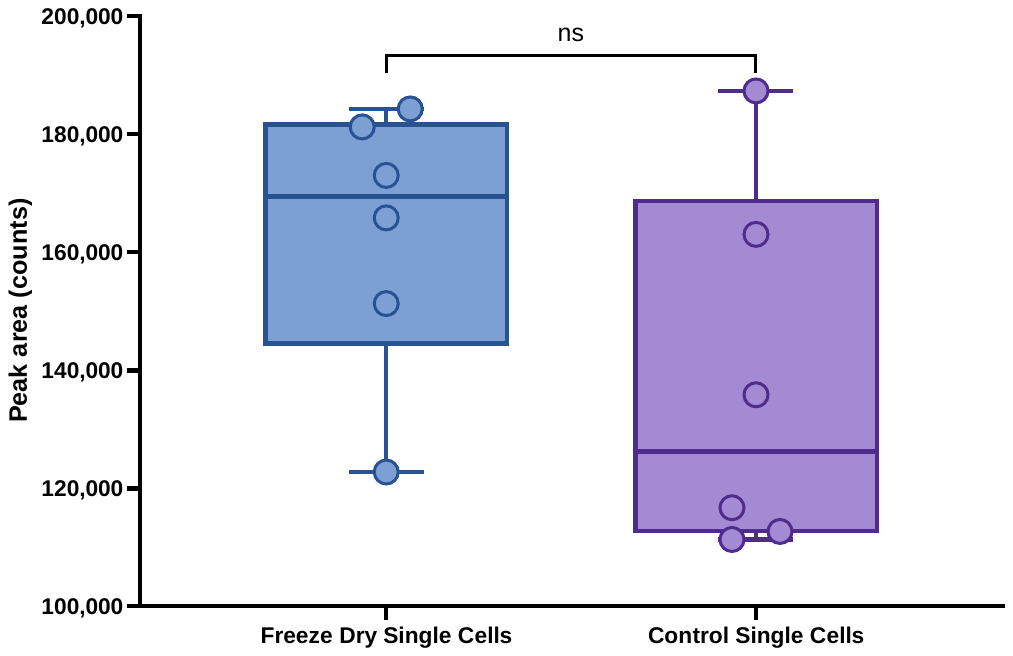

Figure S3. Median peak area of untargeted analysis of freeze dried (blue) and control (purple) single cells analysed using HILIC Q Exactive Plus method.

Table S4. List of analytes from a range of metabolite classes and their formulae monitored for method development.

| Compound | Molecular formula | Class |
| --- | --- | --- |
| Acetyl CoA | C_23_H_38_N_7_O_17_P_3_S | Coenzyme |
| Adenine (Ade) | C_5_H_5_N_5_ | Nucleobase |
| Aspartic acid | C_4_H_7_NO_4_ | Amino Acid |
| Coenzyme A (CoA) | C_21_H_36_N_7_O_16_P_3_S | Coenzyme |
| Glutathione / GSH | C_10_H_17_N_3_O_6_S | Peptide |
| Oleic acid / OA / 18:1 | C_18_H_34_O_2_ | Fatty acid |
| Thymidine / Thd | C_10_H_14_N_2_O_5_ | Nucleoside |
| Tryptophan / Trp | C_11_H_12_N_2_O_2_ | Amino acid |
| Uracil / U | C_4_H_4_N_2_O_2_ | Nucleobase |
| Uridine / Uri | C_9_H_12_N_2_O_6_ | Nucleoside |

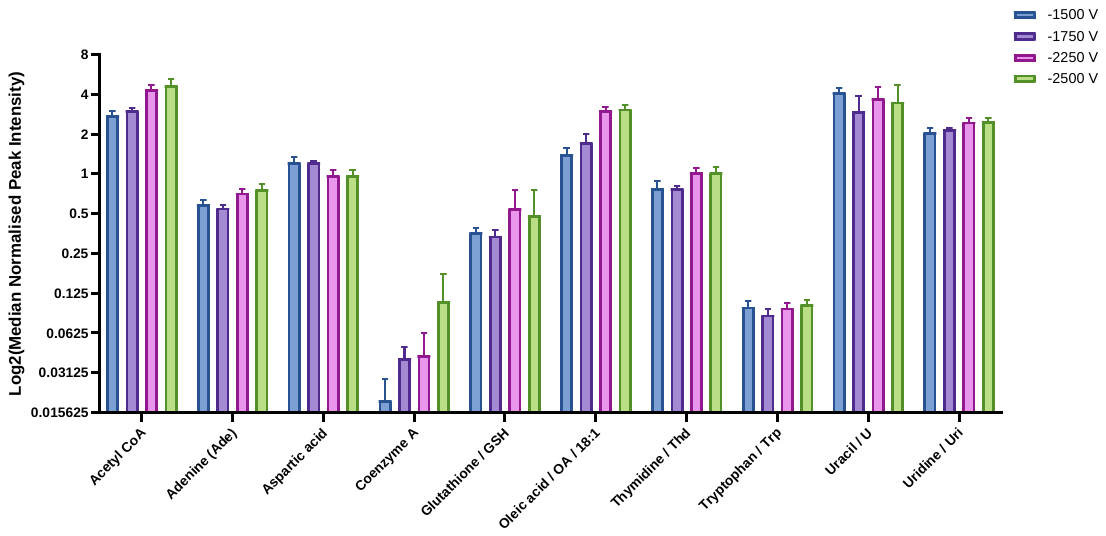

Figure S4. Effect of voltage on metabolites in Table S4 separated and detected using microflow IP method. N = 3.

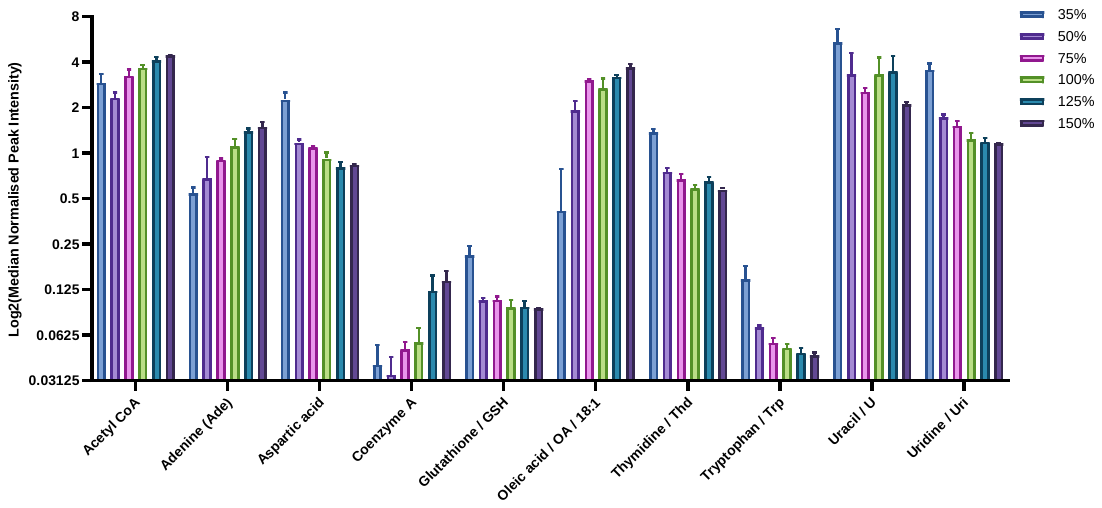

Figure S5. Effect of radio frequency (RF) percentage on metabolites in Table S4 separated and detected using microflow IP method. N = 2.

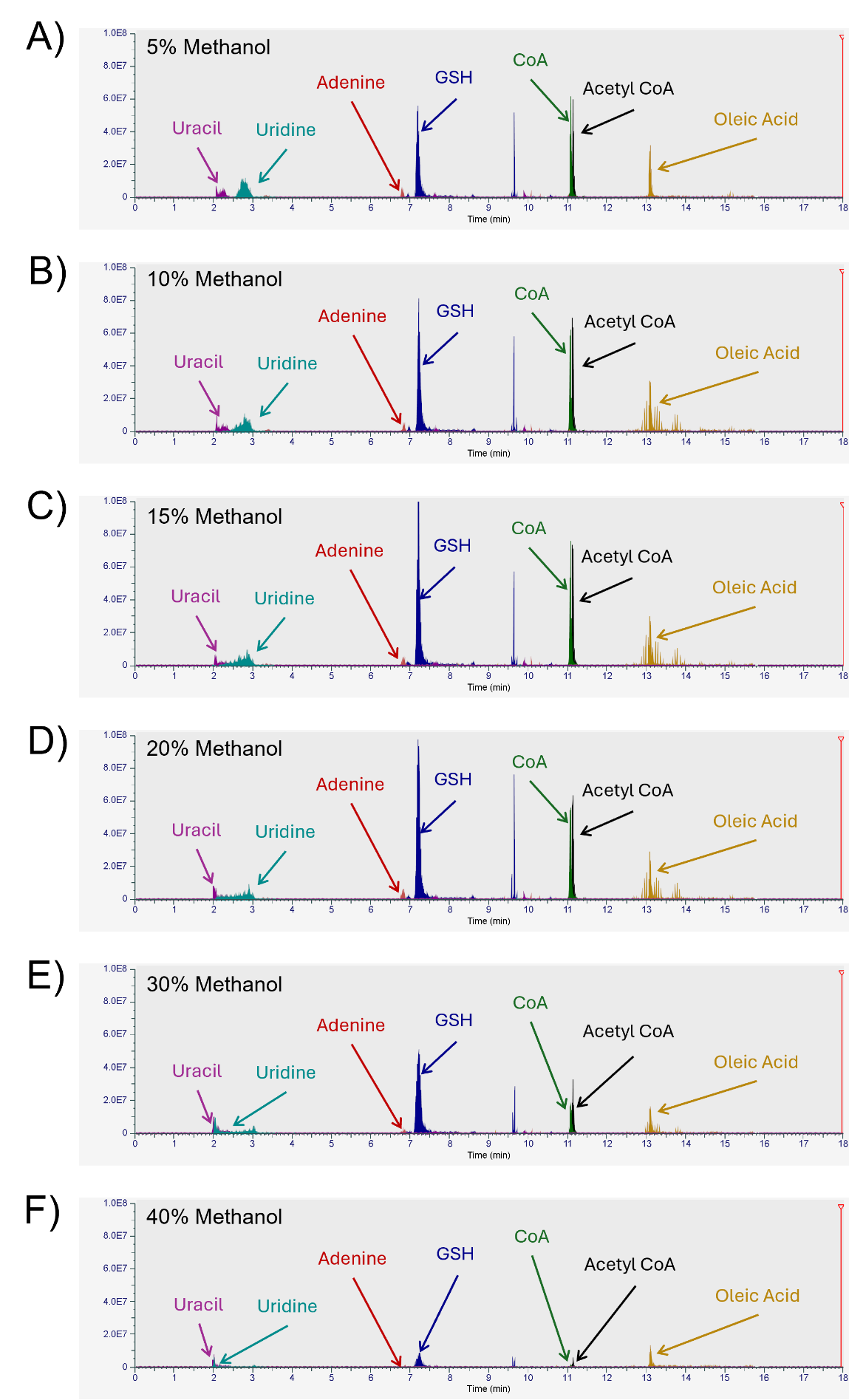

Figure S6. Extracted ion chromatograms displaying the effect of reconstitution solvent composition on peak shape and intensity. A) 5% methanol in water, B) 10% methanol in water, C) 15% methanol in water, D) 20% methanol in water, E) 30% methanol in water, and F) 40% methanol in water. Pink = uracil, light blue = uridine, red = adenine, blue = glutathione (GSH), green = coenzyme A (CoA), black = acetyl CoA, and yellow = oleic acid. The intensity of all chromatograms was normalised to 1e8 AU.

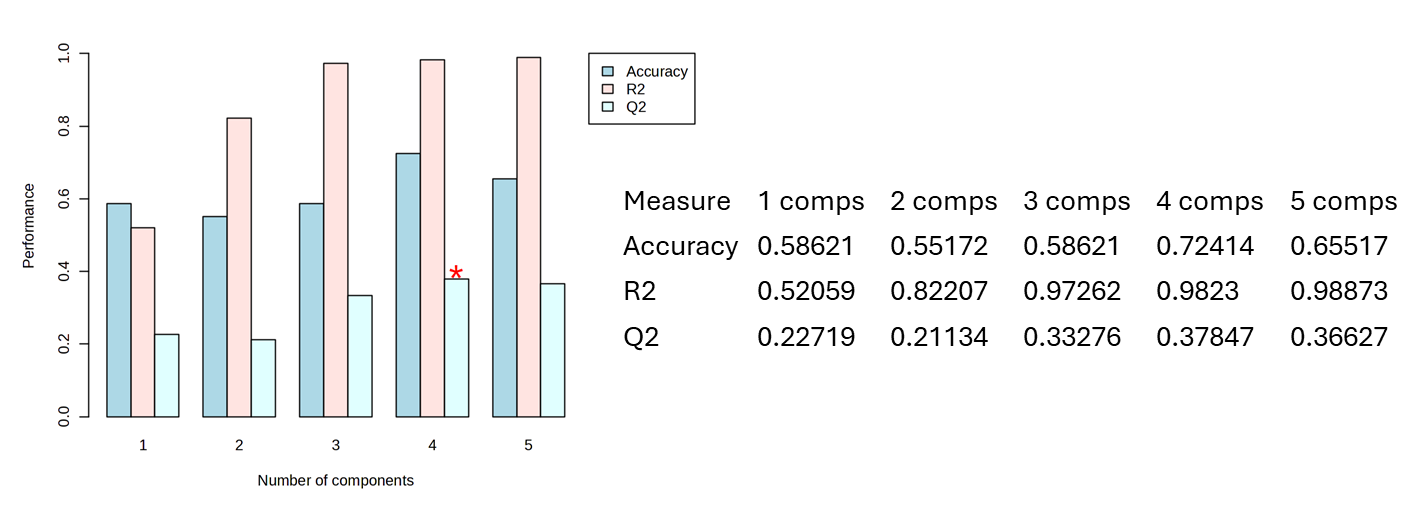

Figure S7. Leave One Out Cross validation (LOOCV) of five components for PLS-DA of IP whole cell clustering of uninfected bystander, BCG infected and unexposed control cell groups.

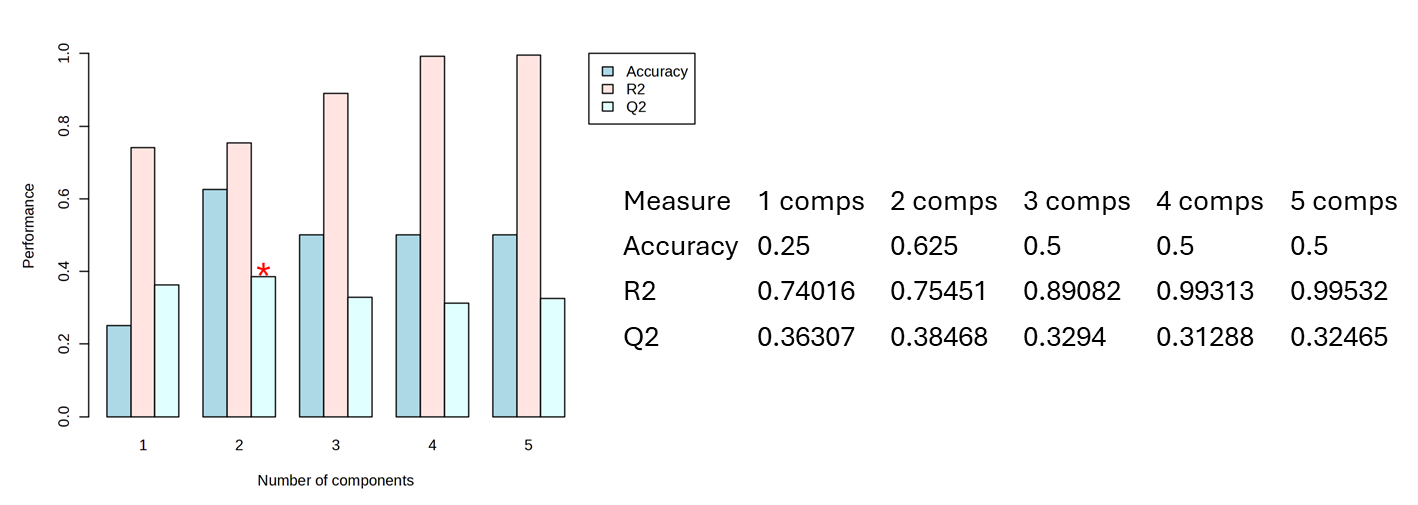

Figure S8. LOOCV of five components for PLS-DA of HILIC whole cell clustering of uninfected bystander, BCG infected and unexposed control cell groups.

Table S5. Pathway enrichment analysis of BCG infected vs unexposed control single cells using HILIC Excedion Pro method.

| Pathway | Total Pathway Compounds | Hits | % Pathway Hit | Raw p | Holm p | FDR |
| --- | --- | --- | --- | --- | --- | --- |
| Arginine biosynthesis | 14 | 6 | 43 | 4.61E-03 | 0.1752 | 0.0526 |
| Nitrogen metabolism | 6 | 3 | 50 | 0.00526 | 0.1945 | 0.0526 |
| Glyoxylate and dicarboxylate metabolism | 32 | 4 | 13 | 0.05062 | 1 | 0.2434 |
| Cysteine and methionine metabolism | 33 | 4 | 12 | 0.06388 | 1 | 0.2555 |
| Pentose phosphate pathway | 22 | 3 | 14 | 0.07121 | 1 | 0.2589 |
| Glycine, serine and threonine metabolism | 32 | 6 | 19 | 0.09648 | 1 | 0.2934 |
| Arginine and proline metabolism | 35 | 18 | 51 | 0.10061 | 1 | 0.2934 |
| Butanoate metabolism | 15 | 4 | 27 | 0.10271 | 1 | 0.2934 |
| Alanine, aspartate and glutamate metabolism | 28 | 10 | 36 | 0.11398 | 1 | 0.304 |
| One carbon pool by folate | 26 | 6 | 23 | 0.15208 | 1 | 0.3714 |
| Terpenoid backbone biosynthesis | 18 | 4 | 22 | 0.16402 | 1 | 0.3714 |
| Purine metabolism | 70 | 17 | 24 | 0.16712 | 1 | 0.3714 |
| Phosphonate and phosphinate metabolism | 6 | 2 | 33 | 0.23667 | 1 | 0.4508 |
| Pyrimidine metabolism | 39 | 16 | 41 | 0.35335 | 1 | 0.6425 |
| Propanoate metabolism | 22 | 5 | 23 | 0.40783 | 1 | 0.6769 |
| D-Amino acid metabolism | 14 | 10 | 71 | 0.42101 | 1 | 0.6769 |
| Phenylalanine metabolism | 8 | 2 | 25 | 0.44978 | 1 | 0.6769 |
| Glycolysis / Gluconeogenesis | 24 | 3 | 13 | 0.47385 | 1 | 0.6769 |
| Pyruvate metabolism | 22 | 3 | 14 | 0.47385 | 1 | 0.6769 |
| Sulfur metabolism | 8 | 2 | 25 | 0.54842 | 1 | 0.7517 |
| Phenylalanine, tyrosine and tryptophan biosynthesis | 4 | 3 | 75 | 0.57228 | 1 | 0.7517 |
| Ubiquinone and other terpenoid-quinone biosynthesis | 20 | 8 | 40 | 0.65992 | 1 | 0.7625 |
| beta-Alanine metabolism | 21 | 7 | 33 | 0.69659 | 1 | 0.7625 |
| Citrate cycle (TCA cycle) | 20 | 5 | 25 | 0.70881 | 1 | 0.7625 |
| Valine, leucine and isoleucine degradation | 40 | 4 | 10 | 0.71628 | 1 | 0.7625 |
| Valine, leucine and isoleucine biosynthesis | 8 | 2 | 25 | 0.73333 | 1 | 0.7625 |
| Pantothenate and CoA biosynthesis | 20 | 10 | 50 | 0.77361 | 1 | 0.7736 |

Table S6. Pathway enrichment analysis of BCG infected vs uninfected bystander single cells using HILIC Excedion Pro method.

| Pathway | Total Pathway Compounds | Hits | % Pathway Hit | Raw p | Holm p | FDR |
| --- | --- | --- | --- | --- | --- | --- |
| Pantothenate and CoA biosynthesis | 20 | 11 | 55 | 3.30E-02 | 1 | 0.4343 |
| Pyrimidine metabolism | 39 | 14 | 36 | 0.04214 | 1 | 0.4343 |
| Butanoate metabolism | 15 | 3 | 20 | 0.07768 | 1 | 0.4946 |
| Cysteine and methionine metabolism | 33 | 5 | 15 | 0.08103 | 1 | 0.4946 |
| beta-Alanine metabolism | 21 | 9 | 43 | 0.13275 | 1 | 0.4946 |
| Pentose phosphate pathway | 22 | 3 | 14 | 0.13326 | 1 | 0.4946 |
| Glycine, serine and threonine metabolism | 32 | 5 | 16 | 0.13949 | 1 | 0.4946 |
| Alanine, aspartate and glutamate metabolism | 28 | 8 | 29 | 0.18302 | 1 | 0.5491 |
| One carbon pool by folate | 26 | 7 | 27 | 0.25617 | 1 | 0.7136 |
| Citrate cycle (TCA cycle) | 20 | 4 | 20 | 0.29361 | 1 | 0.7199 |
| Glyoxylate and dicarboxylate metabolism | 32 | 4 | 13 | 0.29534 | 1 | 0.7199 |
| Pyruvate metabolism | 22 | 3 | 14 | 0.37894 | 1 | 0.7559 |
| Glycolysis / Gluconeogenesis | 24 | 3 | 13 | 0.37894 | 1 | 0.7559 |
| Terpenoid backbone biosynthesis | 18 | 6 | 33 | 0.42641 | 1 | 0.7559 |
| Arginine and proline metabolism | 35 | 18 | 51 | 0.4706 | 1 | 0.7777 |
| Arginine biosynthesis | 14 | 6 | 43 | 0.51761 | 1 | 0.8075 |
| Glutathione metabolism | 28 | 3 | 11 | 0.63024 | 1 | 0.8914 |
| D-Amino acid metabolism | 14 | 11 | 79 | 0.63996 | 1 | 0.8914 |
| Valine, leucine and isoleucine biosynthesis | 8 | 2 | 25 | 0.68771 | 1 | 0.8935 |
| Phenylalanine metabolism | 8 | 2 | 25 | 0.69348 | 1 | 0.8935 |
| Propanoate metabolism | 22 | 5 | 23 | 0.71448 | 1 | 0.8935 |
| Phenylalanine, tyrosine and tryptophan biosynthesis | 4 | 3 | 75 | 0.75664 | 1 | 0.8935 |
| Sulfur metabolism | 8 | 2 | 25 | 0.87933 | 1 | 0.9712 |
| Ubiquinone and other terpenoid-quinone biosynthesis | 20 | 7 | 35 | 0.91386 | 1 | 0.9712 |
| Purine metabolism | 70 | 17 | 24 | 0.92685 | 1 | 0.9712 |
| Nitrogen metabolism | 6 | 3 | 50 | 0.9844 | 1 | 0.9844 |

Table S7. Pathway enrichment analysis of unexposed control vs uninfected bystander single cells using HILIC Excedion Pro method.

| Pathway | Total Pathway Compounds | Hits | % Pathway Hit | Raw p | Holm p | FDR |
| --- | --- | --- | --- | --- | --- | --- |
| Nitrogen metabolism | 6 | 3 | 50 | 1.28E-02 | 0.4736 | 0.128 |
| beta-Alanine metabolism | 21 | 9 | 43 | 0.01856 | 0.6683 | 0.1485 |
| Butanoate metabolism | 15 | 3 | 20 | 0.03994 | 1 | 0.2471 |
| Pyrimidine metabolism | 39 | 13 | 33 | 0.06232 | 1 | 0.262 |
| Arginine biosynthesis | 14 | 5 | 36 | 0.0655 | 1 | 0.262 |
| Arginine and proline metabolism | 35 | 18 | 51 | 0.08455 | 1 | 0.3075 |
| One carbon pool by folate | 26 | 5 | 19 | 0.10035 | 1 | 0.3345 |
| Pantothenate and CoA biosynthesis | 20 | 10 | 50 | 0.10906 | 1 | 0.3356 |
| Glyoxylate and dicarboxylate metabolism | 32 | 4 | 13 | 0.13702 | 1 | 0.3799 |
| Purine metabolism | 70 | 18 | 26 | 0.14864 | 1 | 0.3799 |
| Glycine, serine and threonine metabolism | 32 | 6 | 19 | 0.15196 | 1 | 0.3799 |
| Propanoate metabolism | 22 | 6 | 27 | 0.16487 | 1 | 0.3879 |
| D-Amino acid metabolism | 14 | 10 | 71 | 0.25497 | 1 | 0.5666 |
| Alanine, aspartate and glutamate metabolism | 28 | 8 | 29 | 0.30807 | 1 | 0.6173 |
| Sulfur metabolism | 8 | 2 | 25 | 0.30865 | 1 | 0.6173 |
| Ubiquinone and other terpenoid-quinone biosynthesis | 20 | 7 | 35 | 0.36223 | 1 | 0.69 |
| Valine, leucine and isoleucine biosynthesis | 8 | 2 | 25 | 0.55032 | 1 | 0.9571 |
| Pentose phosphate pathway | 22 | 3 | 14 | 0.64765 | 1 | 0.974 |
| Terpenoid backbone biosynthesis | 18 | 5 | 28 | 0.77613 | 1 | 0.974 |
| Valine, leucine and isoleucine degradation | 40 | 4 | 10 | 0.77717 | 1 | 0.974 |
| Phenylalanine, tyrosine and tryptophan biosynthesis | 4 | 3 | 75 | 0.89484 | 1 | 0.974 |
| Glycolysis / Gluconeogenesis | 24 | 3 | 13 | 0.90958 | 1 | 0.974 |
| Pyruvate metabolism | 22 | 3 | 14 | 0.90958 | 1 | 0.974 |
| Citrate cycle (TCA cycle) | 20 | 4 | 20 | 0.92886 | 1 | 0.974 |
| Phenylalanine metabolism | 8 | 2 | 25 | 0.93656 | 1 | 0.974 |

Table S8. Pathway enrichment analysis of unexposed control vs uninfected bystander single cells using IP Excedion Pro method.

| Pathway | Total Pathway Compounds | Hits | % Pathway Hit | Raw p | Holm p | FDR |
| --- | --- | --- | --- | --- | --- | --- |
| Purine metabolism | 70 | 43 | 61 | 4.56E-05 | 0.0025 | 0.0025 |
| D-Amino acid metabolism | 14 | 13 | 93 | 0.00028 | 0.0152 | 0.0052 |
| Glutathione metabolism | 28 | 3 | 11 | 0.00029 | 0.0152 | 0.0052 |
| Arginine biosynthesis | 14 | 10 | 71 | 0.00062 | 0.0323 | 0.0086 |
| Glycine, serine and threonine metabolism | 32 | 15 | 47 | 0.00392 | 0.1962 | 0.036 |
| One carbon pool by folate | 26 | 16 | 62 | 0.00623 | 0.3054 | 0.0458 |
| Nitrogen metabolism | 6 | 3 | 50 | 0.00667 | 0.3199 | 0.0458 |
| Pentose phosphate pathway | 22 | 4 | 18 | 0.01046 | 0.4916 | 0.0639 |
| Lipoic acid metabolism | 28 | 4 | 14 | 0.01195 | 0.5497 | 0.0657 |
| Amino sugar and nucleotide sugar metabolism | 31 | 8 | 26 | 0.02401 | 1 | 0.1101 |
| Pyrimidine metabolism | 39 | 7 | 18 | 0.02703 | 1 | 0.1144 |
| Glyoxylate and dicarboxylate metabolism | 32 | 9 | 28 | 0.0347 | 1 | 0.1199 |
| Biosynthesis of various nucleotide sugars | 24 | 4 | 17 | 0.03927 | 1 | 0.1199 |
| Valine, leucine and isoleucine biosynthesis | 8 | 6 | 75 | 0.041 | 1 | 0.1199 |
| Folate biosynthesis | 26 | 3 | 12 | 0.04141 | 1 | 0.1199 |
| Valine, leucine and isoleucine degradation | 40 | 6 | 15 | 0.04308 | 1 | 0.1199 |
| Taurine and hypotaurine metabolism | 8 | 2 | 25 | 0.0436 | 1 | 0.1199 |
| Arginine and proline metabolism | 35 | 20 | 57 | 0.04581 | 1 | 0.12 |
| Alanine, aspartate and glutamate metabolism | 28 | 21 | 75 | 0.06131 | 1 | 0.1533 |
| Pantothenate and CoA biosynthesis | 20 | 13 | 65 | 0.07302 | 1 | 0.1746 |
| Porphyrin metabolism | 31 | 11 | 35 | 0.08209 | 1 | 0.1881 |
| Propanoate metabolism | 22 | 5 | 23 | 0.12854 | 1 | 0.2719 |
| Fructose and mannose metabolism | 21 | 13 | 62 | 0.16396 | 1 | 0.3032 |
| Phenylalanine, tyrosine and tryptophan biosynthesis | 4 | 4 | 100 | 0.1655 | 1 | 0.3032 |
| Histidine metabolism | 16 | 10 | 63 | 0.1709 | 1 | 0.3032 |
| Pyruvate metabolism | 22 | 8 | 36 | 0.25517 | 1 | 0.4253 |
| Galactose metabolism | 26 | 3 | 12 | 0.26592 | 1 | 0.4302 |
| Butanoate metabolism | 15 | 4 | 27 | 0.3321 | 1 | 0.5219 |
| Cysteine and methionine metabolism | 33 | 16 | 48 | 0.37342 | 1 | 0.5705 |
| beta-Alanine metabolism | 21 | 10 | 48 | 0.41675 | 1 | 0.6189 |
| Ubiquinone and other terpenoid-quinone biosynthesis | 20 | 13 | 65 | 0.42759 | 1 | 0.6189 |
| Phenylalanine metabolism | 8 | 7 | 88 | 0.47436 | 1 | 0.6471 |
| Sulfur metabolism | 8 | 2 | 25 | 0.48239 | 1 | 0.6471 |
| Citrate cycle (TCA cycle) | 20 | 6 | 30 | 0.52833 | 1 | 0.6919 |
| Arachidonic acid metabolism | 44 | 19 | 43 | 0.60582 | 1 | 0.7573 |
| Glycolysis / Gluconeogenesis | 24 | 8 | 33 | 0.72192 | 1 | 0.879 |
| Vitamin B6 metabolism | 9 | 8 | 89 | 0.79692 | 1 | 0.9131 |
| Nicotinate and nicotinamide metabolism | 15 | 2 | 13 | 0.94443 | 1 | 0.9515 |
| Tyrosine metabolism | 42 | 38 | 90 | 0.95146 | 1 | 0.9515 |

Table S9. Pathway enrichment analysis of BCG infected vs unexposed control single cells using IP Excedion Pro method.

| Pathway | Total Pathway Compounds | Hits | % Pathway Hit | Raw p | Holm p | FDR |
| --- | --- | --- | --- | --- | --- | --- |
| Purine metabolism | 70 | 54 | 77 | 6.38E-05 | 0.0037 | 0.0037 |
| Arginine biosynthesis | 14 | 11 | 79 | 0.00073 | 0.0415 | 0.0211 |
| Pentose phosphate pathway | 22 | 6 | 27 | 0.00167 | 0.0934 | 0.0322 |
| Glutathione metabolism | 28 | 7 | 25 | 0.00335 | 0.1845 | 0.0486 |
| Tryptophan metabolism | 40 | 4 | 10 | 0.0085 | 0.4589 | 0.0854 |
| D-Amino acid metabolism | 14 | 14 | 100 | 0.00883 | 0.468 | 0.0854 |
| Glycine, serine and threonine metabolism | 32 | 21 | 66 | 0.01766 | 0.9185 | 0.1464 |
| One carbon pool by folate | 26 | 20 | 77 | 0.02031 | 1 | 0.1472 |
| Inositol phosphate metabolism | 32 | 4 | 13 | 0.05518 | 1 | 0.3556 |
| Biosynthesis of various nucleotide sugars | 24 | 4 | 17 | 0.07665 | 1 | 0.3811 |
| Nitrogen metabolism | 6 | 3 | 50 | 0.09745 | 1 | 0.3811 |
| Lipoic acid metabolism | 28 | 6 | 21 | 0.10207 | 1 | 0.3811 |
| Histidine metabolism | 16 | 13 | 81 | 0.10328 | 1 | 0.3811 |
| Propanoate metabolism | 22 | 7 | 32 | 0.10973 | 1 | 0.3811 |
| Arginine and proline metabolism | 35 | 24 | 69 | 0.1121 | 1 | 0.3811 |
| Amino sugar and nucleotide sugar metabolism | 31 | 8 | 26 | 0.11304 | 1 | 0.3811 |
| Cysteine and methionine metabolism | 33 | 24 | 73 | 0.11984 | 1 | 0.3811 |
| Pyrimidine metabolism | 39 | 9 | 23 | 0.12026 | 1 | 0.3811 |
| Sulfur metabolism | 8 | 3 | 38 | 0.14238 | 1 | 0.3811 |
| Glycolysis / Gluconeogenesis | 24 | 9 | 38 | 0.19111 | 1 | 0.4573 |
| Fructose and mannose metabolism | 21 | 13 | 62 | 0.20388 | 1 | 0.4573 |
| Glyoxylate and dicarboxylate metabolism | 32 | 11 | 34 | 0.2156 | 1 | 0.4573 |
| Valine, leucine and isoleucine biosynthesis | 8 | 6 | 75 | 0.21856 | 1 | 0.4573 |
| beta-Alanine metabolism | 21 | 14 | 67 | 0.22779 | 1 | 0.4573 |
| Valine, leucine and isoleucine degradation | 40 | 9 | 23 | 0.22866 | 1 | 0.4573 |
| Galactose metabolism | 26 | 3 | 12 | 0.24123 | 1 | 0.4664 |
| Lysine degradation | 29 | 3 | 10 | 0.27438 | 1 | 0.5134 |
| Glycerolipid metabolism | 16 | 2 | 13 | 0.31081 | 1 | 0.5463 |
| Alanine, aspartate and glutamate metabolism | 28 | 25 | 89 | 0.32644 | 1 | 0.5569 |
| Pyruvate metabolism | 22 | 9 | 41 | 0.33867 | 1 | 0.5612 |
| Citrate cycle (TCA cycle) | 20 | 8 | 40 | 0.36871 | 1 | 0.5886 |
| Ubiquinone and other terpenoid-quinone biosynthesis | 20 | 15 | 75 | 0.38388 | 1 | 0.5886 |
| Pantothenate and CoA biosynthesis | 20 | 16 | 80 | 0.39581 | 1 | 0.5886 |
| Taurine and hypotaurine metabolism | 8 | 3 | 38 | 0.40623 | 1 | 0.589 |
| Folate biosynthesis | 26 | 3 | 12 | 0.44465 | 1 | 0.6284 |
| Butanoate metabolism | 15 | 6 | 40 | 0.45619 | 1 | 0.6284 |
| Vitamin B6 metabolism | 9 | 9 | 100 | 0.47956 | 1 | 0.6284 |
| Porphyrin metabolism | 31 | 16 | 52 | 0.51305 | 1 | 0.6284 |
| Terpenoid backbone biosynthesis | 18 | 2 | 11 | 0.69047 | 1 | 0.7701 |
| Arachidonic acid metabolism | 44 | 38 | 86 | 0.78664 | 1 | 0.8554 |
| Tyrosine metabolism | 42 | 39 | 93 | 0.79637 | 1 | 0.8554 |
| Phenylalanine, tyrosine and tryptophan biosynthesis | 4 | 4 | 100 | 0.85712 | 1 | 0.9039 |
| Nicotinate and nicotinamide metabolism | 15 | 2 | 13 | 0.91748 | 1 | 0.9336 |
| Phenylalanine metabolism | 8 | 8 | 100 | 0.94944 | 1 | 0.9494 |

Table S10. Pathway enrichment analysis of BCG infected vs uninfected bystander single cells using IP Excedion Pro method.

| Pathway | Total Pathway Compounds | Hits | % Pathway Hit | Raw p | Holm p | FDR |
| --- | --- | --- | --- | --- | --- | --- |
| Lipoic acid metabolism | 28 | 6 | 21 | 2.73E-02 | 1 | 0.9952 |
| Purine metabolism | 70 | 46 | 66 | 0.13825 | 1 | 0.9952 |
| Tryptophan metabolism | 40 | 4 | 10 | 0.16444 | 1 | 0.9952 |
| Lysine degradation | 29 | 3 | 10 | 0.18754 | 1 | 0.9952 |
| One carbon pool by folate | 26 | 16 | 62 | 0.19346 | 1 | 0.9952 |
| Terpenoid backbone biosynthesis | 18 | 2 | 11 | 0.3563 | 1 | 0.9952 |
| Phenylalanine, tyrosine and tryptophan biosynthesis | 4 | 4 | 100 | 0.41153 | 1 | 0.9952 |
| Glutathione metabolism | 28 | 5 | 18 | 0.43839 | 1 | 0.9952 |
| Vitamin B6 metabolism | 9 | 9 | 100 | 0.45566 | 1 | 0.9952 |
| Taurine and hypotaurine metabolism | 8 | 2 | 25 | 0.59327 | 1 | 0.9952 |
| Ubiquinone and other terpenoid-quinone biosynthesis | 20 | 15 | 75 | 0.60333 | 1 | 0.9952 |
| Glyoxylate and dicarboxylate metabolism | 32 | 10 | 31 | 0.63605 | 1 | 0.9952 |
| Pantothenate and CoA biosynthesis | 20 | 12 | 60 | 0.66774 | 1 | 0.9952 |
| Glycine, serine and threonine metabolism | 32 | 16 | 50 | 0.68466 | 1 | 0.9952 |
| Pyrimidine metabolism | 39 | 6 | 15 | 0.698 | 1 | 0.9952 |
| Cysteine and methionine metabolism | 33 | 20 | 61 | 0.732 | 1 | 0.9952 |
| Pentose phosphate pathway | 22 | 5 | 23 | 0.74308 | 1 | 0.9952 |
| Inositol phosphate metabolism | 32 | 4 | 13 | 0.74745 | 1 | 0.9952 |
| Porphyrin metabolism | 31 | 14 | 45 | 0.75907 | 1 | 0.9952 |
| Amino sugar and nucleotide sugar metabolism | 31 | 8 | 26 | 0.76897 | 1 | 0.9952 |
| beta-Alanine metabolism | 21 | 11 | 52 | 0.77178 | 1 | 0.9952 |
| D-Amino acid metabolism | 14 | 13 | 93 | 0.77815 | 1 | 0.9952 |
| Valine, leucine and isoleucine degradation | 40 | 7 | 18 | 0.77889 | 1 | 0.9952 |
| Phenylalanine metabolism | 8 | 8 | 100 | 0.81584 | 1 | 0.9952 |
| Histidine metabolism | 16 | 10 | 63 | 0.82375 | 1 | 0.9952 |
| Nitrogen metabolism | 6 | 2 | 33 | 0.8307 | 1 | 0.9952 |
| Glycolysis / Gluconeogenesis | 24 | 9 | 38 | 0.84649 | 1 | 0.9952 |
| Pyruvate metabolism | 22 | 9 | 41 | 0.85569 | 1 | 0.9952 |
| Tyrosine metabolism | 42 | 38 | 90 | 0.87087 | 1 | 0.9952 |
| Arginine and proline metabolism | 35 | 22 | 63 | 0.88736 | 1 | 0.9952 |
| Valine, leucine and isoleucine biosynthesis | 8 | 5 | 63 | 0.89097 | 1 | 0.9952 |
| Galactose metabolism | 26 | 3 | 12 | 0.89746 | 1 | 0.9952 |
| Propanoate metabolism | 22 | 5 | 23 | 0.93051 | 1 | 0.9952 |
| Biosynthesis of various nucleotide sugars | 24 | 4 | 17 | 0.93201 | 1 | 0.9952 |
| Fructose and mannose metabolism | 21 | 13 | 62 | 0.93658 | 1 | 0.9952 |
| Alanine, aspartate and glutamate metabolism | 28 | 21 | 75 | 0.93929 | 1 | 0.9952 |
| Citrate cycle (TCA cycle) | 20 | 8 | 40 | 0.95192 | 1 | 0.9952 |
| Sulfur metabolism | 8 | 2 | 25 | 0.95661 | 1 | 0.9952 |
| Arginine biosynthesis | 14 | 10 | 71 | 0.9569 | 1 | 0.9952 |
| Nicotinate and nicotinamide metabolism | 15 | 2 | 13 | 0.96032 | 1 | 0.9952 |
| Arachidonic acid metabolism | 44 | 38 | 86 | 0.98926 | 1 | 0.9963 |
| Butanoate metabolism | 15 | 5 | 33 | 0.99634 | 1 | 0.9963 |

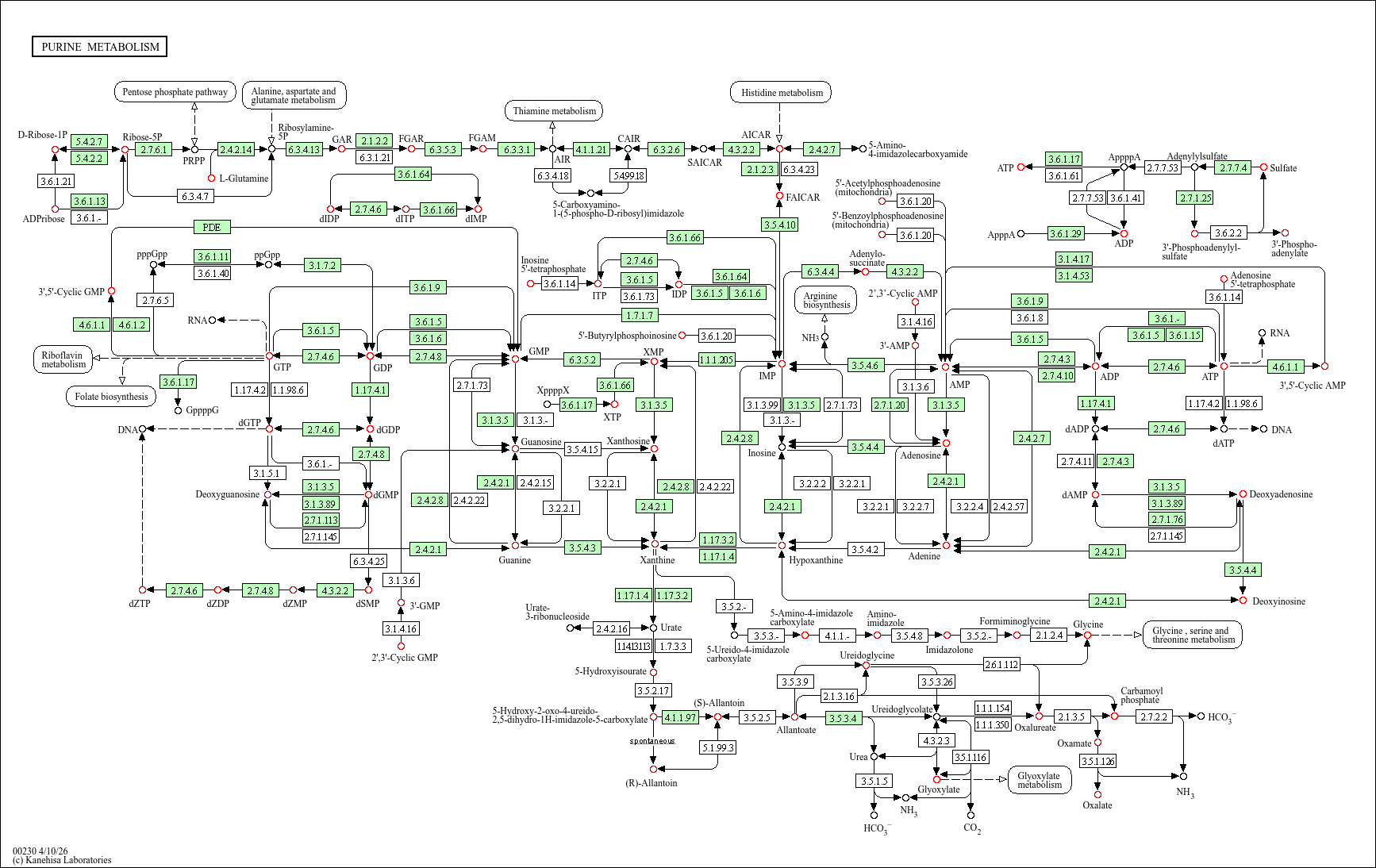

Figure S9. KEGG purine metabolism pathway in *Homo sapiens*. Detected analytes are highlighted in red. The boxes represent a gene product where those highlighted in green are specific to *H. sapiens*. <https://www.kegg.jp/pathway/hsa00230>

Table S11. Pairwise F test p values for measured Log10 and median normalised peak areas of glycine and ATP.

|  | Bystander vs Control  F test p | Control vs Infected  F test p | Bystander vs Infected  F test p |
| --- | --- | --- | --- |
| Glycine | 0.0038 | 0.7648 | 0.0093 |
| ATP | 0.0726 | 0.0421 | 0.0006 |

Table S12. Interquartile range (IQR) and median absolute deviation (MAD) of Log10 and median normalised peak areas of glycine and ATP.

|  | Glycine | | ATP | |
| --- | --- | --- | --- | --- |
|  | IQR (au) | MAD (au) | IQR (au) | MAD (au) |
| Bystander | 0.525 | 0.182 | 2.439 | 0.6126 |
| Control | 2.2545 | 1.0283 | 0.459 | 0.2434 |
| Infected | 1.5692 | 0.6943 | 0.6251 | 0.2393 |

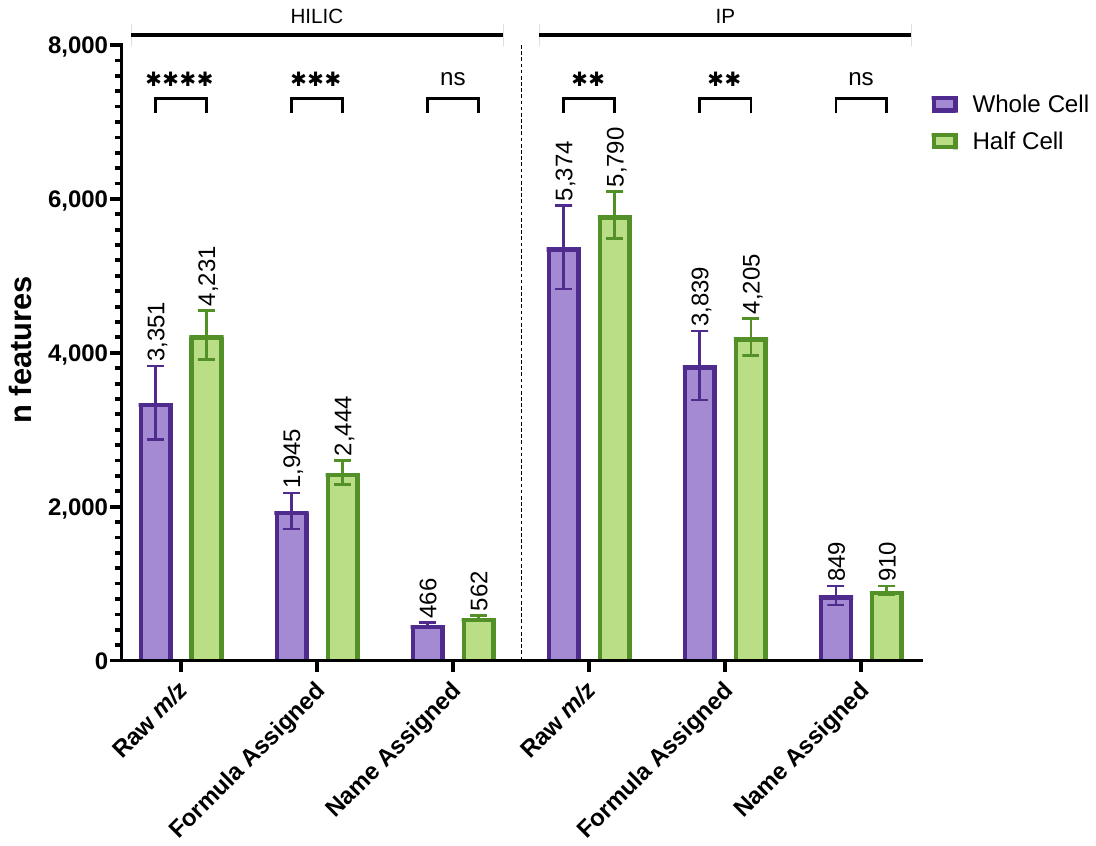

Figure S10. Comparison of number average number of features detected in whole cells (pink) vs half cells (green) analysed on HILIC (left) and/or IP (right) methods. Significance is determined by calculated by a 2-way ANOVA with Holm-Šídák correction for multiple comparisons. HILIC Whole Cell n = 16, IP Whole Cell n = 29, HILIC & IP Split Cell n = 12.

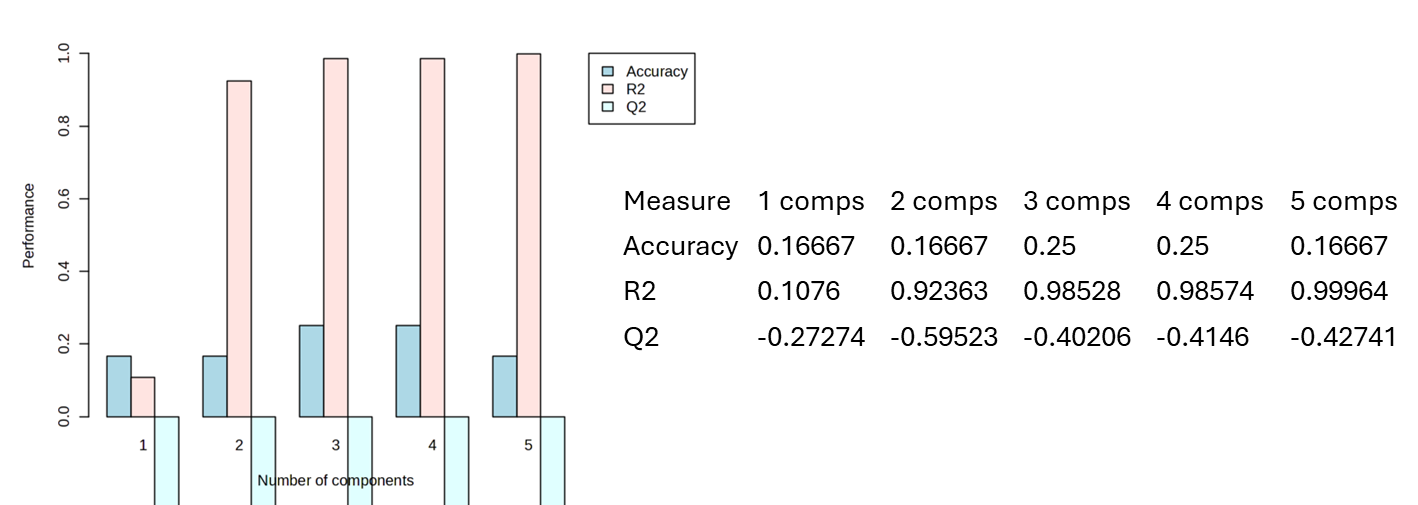

Figure S11. LOOCV of five components for PLS-DA of IP and HILIC clustering of uninfected bystander, BCG infected and unexposed control cell groups.
